# Diverse endogenous retroviruses generate structural variation between human genomes via LTR recombination

**DOI:** 10.1101/382630

**Authors:** Jainy Thomas, Hervé Perron, Cédric Feschotte

## Abstract

Human endogenous retroviruses (HERVs) occupy a substantial fraction of the genome and impact cellular function with both beneficial and deleterious consequences. The vast majority of HERV sequences descend from ancient retroviral families no longer capable of infection or genomic propagation. In fact, most are no longer represented by full-length proviruses but by solitary long terminal repeats (solo LTRs) that arose via non-allelic recombination events between the two LTRs of a proviral insertion. Because LTR-LTR recombination events may occur long after proviral insertion but are challenging to detect in resequencing data, we hypothesize that this mechanism produces an underappreciated amount of genomic variation in the human population. To test this idea, we develop a computational pipeline specifically designed to capture such dimorphic HERV alleles from short-read genome sequencing data. When applied to 279 individuals sequenced as part of the Simons Genome Diversity Project, the pipeline retrieves most of the dimorphic variants previously reported for the HERV-K(HML2) subfamily as well as dozens of additional candidates, including members of the HERV-H and HERV-W families. We experimentally validate several of these candidates, including the first reported instance of an unfixed HERV-W provirus. These data indicate that human proviral content exhibit more extensive interindividual variation than previously recognized. These findings have important implications for our understanding of the contribution of HERVs to human physiology and disease.

## INTRODUCTION

Endogenous retroviruses (ERVs) derive from exogenous retroviruses that inserted in the germline of their host and thereby became vertically inheritable. Full-length (proviral) ERV insertions are comprised of two long terminal repeats (LTRs) flanking an internal region encoding the protein-coding genes necessary for retroviral replication and propagation, including *gag* (group antigens); *pol* (polymerase) and *env* (envelope) (1, 2). ERV sequences are abundant in mammalian genomes, occupying approximately 5 to 10% of the genetic material (3, 4), but virtually each species is unique for its ERV content (5, 6). Indeed, while a fraction of ERVs descend from ancient infections that occurred prior to the emergence of placental mammals, most are derived from independent waves of invasion from diverse viral progenitors that succeeded throughout mammalian evolution (7, 8). Thus, ERVs represent an important source of genomic variation across and within species, including humans. The accumulation of ERV sequences in mammalian genomes has also provided an abundant raw material, both coding and regulatory, occasionally co-opted to foster the emergence of new cellular functions (2, 9–11).

A considerable amount of work has been invested in investigating the pathogenic impact of ERVs. ERVs are prominent insertional mutagens in some species, such as in the mouse where many *de novo* ERV insertions disrupting gene functions have been identified, including tumorigenic insertions (1, 12–14). In contrast, there is no evidence for any *de novo* ERV insertion in humans. However, overexpression of certain human ERV (HERV) families has been associated with a number of disease states, including a variety of cancers, autoimmune, and neurological diseases (15–20). It is becoming increasingly apparent that elevated levels of HERV-derived products, either RNA or proteins, can have direct pathogenic effects (21, 22). However, the genomic mechanisms underlying the differential expression of ERV products in diseased individuals remain obscure. Copy number variation represents a potent mechanism to create inter-individual differences in HERV expression (23), but the extent by which HERV genes vary in copy number across humans and how this variation relates to disease susceptibility remains understudied.

Copy number variation in ERV genes may occur through two primary mechanisms: (i) insertion polymorphisms whereby one allele corresponds to the full provirus while the ancestral allele is completely devoid of the element; (ii) ectopic homologous recombination between the LTRs of the provirus, which results in the deletion of the internal coding sequence, leaving behind a solitary (or solo) LTR (2, 24) (Figure 1 A-C). Thus, one can distinguish three allelic states for ERV insertions: empty, proviral, and solo LTR (25, 26). The process of LTR-LTR recombination has been remarkably efficient in evolution since ~90% of all human ERV (HERV) insertions are currently represented by solo LTRs in the reference genome (27). In theory, the formation of solo LTR from a provirus may occur long after the initial proviral insertion as long as there is sufficient sequence similarity between the two LTRs to promote their recombination. The consequences of this recombination process for the host organism may be significant: not only it removes the entire coding potential of a provirus, but it may also alter the cis-regulatory or transcriptional activity of the LTR (28–32).

**Figure 1.**
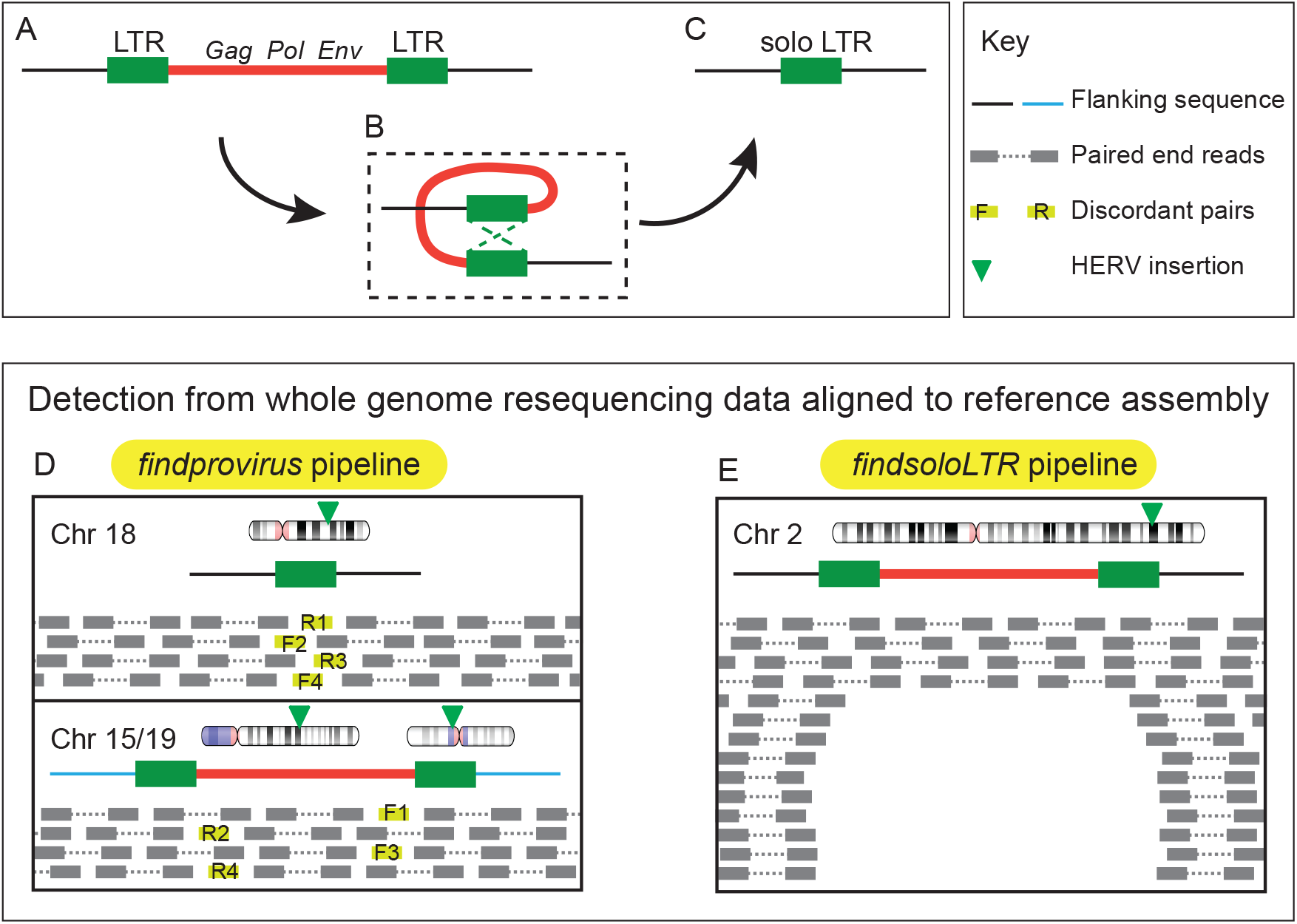
Structure of a provirus and generation of a solo LTR and their detection from whole genome sequence data. Structure of a typical provirus (A) with its internal region (red line) encoding *gag, pol* and *env* genes flanked by two long terminal repeats (LTR). Ectopic recombination occurs between the two LTRs of the provirus (B) leading to the deletion of the internal region along with one LTR, resulting in the formation of a solo LTR (C). Note how the 5’ and 3’ junction sequences between the element and the flanking host DNA (black line), including the target site duplication (not shown), remain the same after recombination. Presence of provirus is identified from whole genome resequencing data aligned to the reference assembly when the reference allele is a solo LTR using the *findprovirus* pipeline (D). The *findprovirus* pipeline infer the presence of provirus from the mates of discordant reads with significant homology to the internal region of the respective HERV family. The *findsoloLTR* pipeline identifies the presence of solo LTR when the reference allele is provirus (E). It infers the presence of solo LTR based on the deviation of read depth across the provirus and across the flank.

Among the diverse assemblage of HERV families in our genome, a single subfamily known as HERV-K(HML2) has been reported to exhibit insertional polymorphism in humans (25–27, 33–44). Thus far, approximately 50 HERV-K(HML2) proviral loci are known to occur as empty (preintegration) and/or solo LTR alleles segregating in the human population (26, 40, 42, 43), but more may be expected to segregate at low frequency (36, 45). These observations are consistent with the notion that HERV-K(HML2) is the most recently active HERV subfamily in the human genome (46–50). To our knowledge, there has been only a single report of another HERV family exhibiting a dimorphic locus: an HERV-H element on chromosome 1 (1q25.3_H3) was shown to exist as proviral and solo LTR alleles in two related individuals (24). Because LTR recombination may in principle took place long after a proviral insertion has reached fixation (51) and possibly recur in multiple individuals, we hypothesized that many more proviral-to-solo HERV variants occur in the human population. We also surmised that this type of dimorphic variants could easily escape detection with current computational pipelines. Indeed, these tools are, by design, geared toward the identification of structural breakpoints distinguishing empty and insertion alleles (26, 52–54). By contrast, proviral and solo LTR allelic variants share the same exact junctions with flanking host DNA, thus making them recalcitrant to detection with tools tailored to map insertional polymorphisms.

Here we introduce a novel computational pipeline specifically geared toward the identification of proviral deletion resulting from LTR recombination events. We apply the pipeline to the analysis of genome sequences from 279 individuals from worldwide populations generated as part of the Simons Genome Diversity Project (SGDP) (55). Our approach identifies most dimorphic HERV-K(HML2) loci previously recognized in other population datasets as well as multiple candidate dimorphic HERV-H and HERV-W loci, several of which we validate experimentally. Our results suggest that LTR recombination is an underappreciated source of structural variation in human genomes generating potentially physiologically significant differences in proviral gene copy numbers between individuals.

## RESULTS

### Strategy for identification of proviral allele when the reference allele is a solo LTR

We developed a pipeline called *findprovirus* to mine whole genome resequencing data to detect a proviral allele of a locus annotated as a solo LTR in the reference genome (Figure 1D). The prediction is that a fraction of the read mates to the reads mapping to the annotated solo LTR should be derived from internal sequences of the provirus allele. When mapped to the reference genome, these events should be identified as discordant read mates mapping elsewhere in the reference genome as they may frequently map to the internal region of non-allelic proviral copies. The pipeline extracts reads mapped to the solo LTR and mates of discordant reads to conduct homology-based searches using the discordant read mates as queries against the consensus sequence of the internal region of the respective provirus as defined in the Repbase database (56) (see also Methods). Presence of at least four reads with significant homology to the internal sequence indicates the presence of a potential allele containing a provirus.

In addition to the principal approach described above, the pipeline employs two alternate methods to detect the presence of a provirus at a locus (Figure S1). First, average read depth at the solo LTR is compared to the average of read depth of all solo LTRs in the same individual genome. If the sequenced individual has at least one provirus allele instead of a solo LTR (as in the reference genome), we predict to see an increase in the number of uniquely mapping reads mapping to the solo LTR. Indeed, reads derived to the 5’ and 3’ LTR of the proviral allele remain more likely to map uniquely to the solo LTR than to other LTRs located elsewhere in the reference genome. This is because gene conversion events frequently homogenize the sequence of proviral LTRs (57, 58). Hence the reads derived from the two LTRs of the provirus will preferentially map to the solo LTR annotated in the reference genome, resulting in an increase in read depth at this LTR relative to other solo LTRs in the genome (Figure S2). Second, a local *de novo* assembly of all reads including mates is performed and failure to assemble a solo LTR allele is interpreted as an indicator of the presence of two proviral alleles at the locus (Figure S1, see Methods). Overall the *findprovirus* pipeline predicts the presence of a proviral allele based primarily on the first approach with results from the two alternate approaches used as secondary indicators.

### Known and new dimorphic HERVs predicted through the *findprovirus* pipeline

The *findprovirus* pipeline was used to identity dimorphic candidates for HERV-K(HML2), (hereafter simply noted as HERV-K), HERV-H, and HERV-W families in a dataset consisting of whole genome sequence data for 279 individuals from the SGDP (55). Solo LTRs annotated in the hg38 reference genome for HERV-K (LTR5_Hs) (n= 553), HERV-H (LTR7) (n= 689) and HERV-W (LTR17) (n= 476) were used as initial queries (see Methods). The pipeline reports the following results: (i) number of discordant reads mapping to the region; (ii) number of informative discordant reads (i.e. their mates have a significant hit with the respective HERV coding sequence); (iii) percentage of reference solo LTR allele aligned to *de novo* assembled contigs from the reads; (iv) ratio of average read depth of the element to the average read depth at all solo LTRs of that individual; (v) average mappability of regions where informative discordant reads are mapped; and (vi) prediction on the presence or absence of the provirus allele. The candidates are then visually inspected using Integrative Genomics Viewer (IGV) for the presence of nested polymorphic transposable element (TE) insertion or presence of internal region of same HERV nearby that could result in false positives. After *in silico* inspection, we identify three strong candidate loci for HERV-K, two for HERV-H, and one for HERV-W (see Table 1, Table S1). Two of the three HERV-K candidates have been previously identified and experimentally validated as dimorphic in prior studies (27, 41, 43) (Table 1). For these two loci, we also identified genomic sequences of the corresponding proviral alleles from the Nucleotide collection (nr/nt) database at National Centre for Biotechnology Information (NCBI) through homology-based searches (see methods) (Table S1). The novel dimorphic candidate that we identified for HERV-K (5q11.2_K3) is predicted to be a provirus in 164 individuals and a maximum of six informative discordant reads are mapped to that locus in an individual (Table S1). However, the low average mappability scores for the solo LTR region where the informative discordant reads are mapped suggest that it is a region prone to ambiguous mapping (Table S1). Further experimental validations will be necessary to confirm this dimorphism. Nonetheless, these results show that our pipeline efficiently retrieves known dimorphic HERV-K elements.

**Table 1.** Dimorphic HERV-K, HERV-H and HERV-W candidates

| HERV name | Coordinate (GRCh38/hg38) | Previously reported |
| --- | --- | --- |
| 1p31.1_K2<sup>#†</sup> | chr1:73129298-73130265 | (43) |
| K111/K105<sup>#</sup> | chrUn_GL000219v1:175210-176178 | (27, 41) |
| 1p31.1_K3<sup>‡</sup> | chr1:75377086-75383458 | (25) |
| 11q22.1_K1<sup>‡</sup> | chr11:101695063-101704528 | (25) |
| 12q14.1_K1<sup>‡</sup> | chr12:58327459-58336915 | (25) |
| 3q13.2_K2<sup>‡</sup> | chr3:113024277-113033435 | (35) |
| 3q27.2_K1 | chr3:185562548-185571727 | (40) |
| 5q33.3_K1<sup>‡</sup> | chr5:156657706-156666885 | (40) |
| 6q14.1_K1<sup>‡</sup> | chr6:77716945-77726366 | (25) |
| 7p22.1_K1 | chr7:4582426-4591897 | (25,35) |
| 5p13.3_K2<sup>‡*</sup> | chr5:30486653-30496098 | This study |
| 4q22.1_H8<sup>†*#</sup> | chr4:91045790-91046151 | This study |
| 5p15.31_H2<sup>†*#</sup> | chr5:7262337-7262742 | This study |
| 11q13.2_H5<sup>‡</sup> | chr11:68633778-68639439 | This study |
| 2q34_H4<sup>‡*</sup> | chr2:209078020-209084376 | This study |
| 2p14_H2<sup>‡</sup> | chr2:64252414-64257646 | This study |
| 13q21.32_H1<sup>‡</sup> | chr13:66141332-66147036 | This study |
| 1q32.2_H3<sup>‡</sup> | chr1:210111090-210116207 | This study |
| 1p32.3_H6<sup>‡</sup> | chr1:54897904-54903584 | This study |
| 11q24.3_H2<sup>‡</sup> | chr11:130753498-130759137 | This study |
| 11p14.3_H1<sup>‡</sup> | chr11:23183934-23189744 | This study |
| 12p12.1_H2<sup>‡</sup> | chr12:25163213-25169508 | This study |
| 13q21.1_H1<sup>‡</sup> | chr13:55578228-55584087 | This study |
| 13q22.3_H1<sup>‡</sup> | chr13:77933223-77939379 | This study |
| 2q36.1_H5<sup>‡</sup> | chr2:224296633-224302363 | This study |
| 2p12_H2<sup>‡</sup> | chr2:75213731-75219537 | This study |
| 3q22.3_H2<sup>‡</sup> | chr3:137595601-137601190 | This study |
| 3p14.3_H1<sup>‡*</sup> | chr3:54634484-54640204 | This study |
| 4q32.3_H5<sup>‡</sup> | chr4:166716125-166722054 | This study |
| 6q23.2_H3<sup>‡</sup> | chr6:131338800-131344564 | This study |
| 6p22.3_H3<sup>‡</sup> | chr6:18754144-18759870 | This study |
| 6p12.2_H1<sup>‡</sup> | chr6:51938241-51944426 | This study |
| 6q16.1_H1<sup>‡</sup> | chr6:93830156-93835749 | This study |
| 18q21.1_W2<sup>†*#</sup> | chr18:50449151-50449914 | This study |

To the best of our knowledge, none of the dimorphic HERV-H and HERV-W candidates identified herein have been reported in the literature. The two HERV-H candidates were flagged by up to 23 and 6 discordant mate reads aligned to the internal sequence of HERV-H in an individual (Table 1, Table S1). The HERV-W candidate, 18q21.1_W2 displayed up to 33 discordant mates aligned to HERV-W internal sequence in a given individual (Figure S2). The *findprovirus* pipeline predicted that 194 of 279 individuals had at least one proviral allele of 18q21.1_W2, suggesting that this is a common allele in the human population (Table S1). To experimentally validate these three candidates (Table 1), we used Polymerase Chain Reaction (PCR) to genotype a panel of individuals from the SGDP predicted to include a mixture of genotypes. Primers were designed in the flanking regions and used as a pair to detect the solo LTR allele or in combination with an internal primer (located in *gag* and/or *env* region) to detect the proviral allele (see Methods). The PCR products were analyzed by gel electrophoresis and their identity was confirmed by Sanger sequencing (Supplementary Data 1). The results validated that each of the three loci exist as proviral and solo LTR alleles in the human population (Figure 2 A-C)(Table 1). In addition, we also identified seven FOSMID clones in the nr/nt database at NCBI supporting the presence of proviral alleles (Table S1). Altogether these data strongly support the dimorphic HERV-H and HERV-W calls made through our *findprovirus* pipeline.

**Figure 2.**
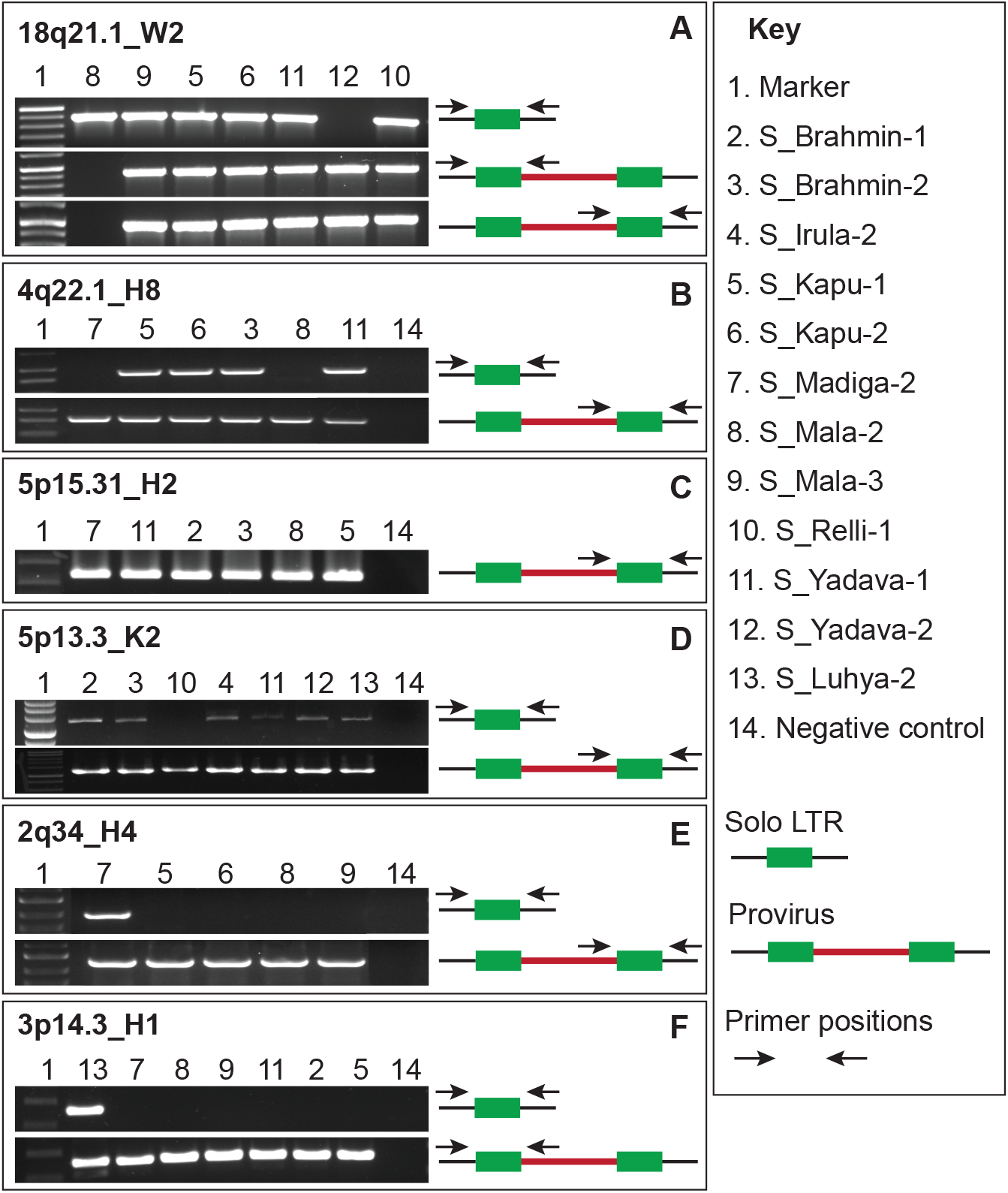
Experimental validation of dimorphic HERV loci. A) PCR amplification of HERV-W solo LTR at the 18q21.1 locus. Primers were designed flanking the solo LTR. PCR amplification of the 18q21.1_W2 provirus with primers designed to the flank and internal *gag* sequence and with primers to the *env* sequence and flank. B) PCR amplification of HERV-H solo LTR at the 4q22.1 locus with primers flanking the solo LTR. PCR amplification of the 4q22.1_H8 provirus with primers designed to the internal *env* sequence and flank. C) PCR amplification of HERV-H provirus at the 5p15.31 locus with primers designed to the internal *env* sequence and flank. D) PCR amplification of HERV-H solo LTR at the at 5p13.3 locus with primers flanking the solo LTR. PCR amplification of the provirus 5p13.3_K2 with primers designed to the internal *env* sequence and flank. E) PCR amplification of HERV-H solo LTR at 2q34 locus with primers flanking the solo LTR. PCR amplification of the provirus 2q34_H4 with primers designed to the internal *env* sequence and flank. F) PCR amplification of HERV-H solo LTR at 3p14.3 locus with primers flanking the solo LTR. PCR amplification of the provirus 3p14.3_H1 with primers designed to the internal *gag* sequence and flank. The DNA samples of various South Asian populations and an African individual used for validation are listed in the key. LTRs are in shown as green boxes, the internal region as a red line, the flanking region as a black line. The primer positions are shown as black arrows.

### Strategy for identification of solo LTR allele when the reference allele is a provirus

We developed a complementary pipeline called *findsoloLTR* to mine whole genome resequencing data to detect a solo LTR allele of a locus annotated as a provirus in the reference genome (Figure 1E). Here the prediction is that an individual with one copy of a proviral allele instead of two will have a decreased number of reads mapping uniquely (mapping quality >=30) to the internal region and an individual with two solo LTR alleles will have even fewer reads mapping uniquely to the internal region of the provirus. The *findsoloLTR* pipeline systematically measures the read depth across the provirus and in the flanking 250-bp regions of the provirus. The pipeline then express the average read depth across the provirus as the percentage of the average read depth across its flanking genomic regions (Figure S3). The candidate locus is considered harboring a solo LTR allele when the calculated read depth ratio across the provirus is lower than 50%. The presence of two solo LTRs alleles is inferred when read depth gets lower than 10% in comparison with the average read depth of the flanking regions (Figure S4).

### Known and new dimorphic HERVs predicted through the *findsoloLTR* pipeline

The *findsoloLTR* pipeline was used to analyze the SGDP data for the presence of solo LTR alleles to a set of sequences annotated as proviruses in the reference genome for HERV-K (n= 23), HERV-H (n= 720) and HERV-W (n= 53). The *findsoloLTR* pipeline reports: (i) mean read depth across the provirus, (ii) mean read depth of the 5’ and 3’ flanks, (iii) percentage of read depth at the provirus to the average of read depth of the flanks and (iv) prediction of the presence of a solo LTR allele. The candidates were visually inspected using IGV to assess whether the decreased read depth ratio was due to a partial deletion instead of the outcome expected for a LTR recombination event which precisely deletes one LTR along with the internal sequence (see Figure S4 for a legitimate candidate). After *in silico* inspection, we retained 12 HERV-K candidates, 67 HERV-H candidates, and no HERV-W candidate (Table 1, Table S2).

In the case of HERV-K, eight of the 12 candidate loci were previously reported to be dimorphic, and some were known to be also insertionally polymorphic, i.e. a pre-integration ‘empty’ allele has also been reported (25, 27, 35, 40, 43) (see Table 1, Table S2). The pipeline predicts four novel HERV-K loci to be dimorphic in the population. For HERV-H, we observe that many of the predicted solo LTR allele occurs at low frequency in the SGDP dataset, being predicted in only a few individuals (Table 1, Table S2). This might be expected if these alleles arose from relatively recent recombination events. Alternatively, they may represent false positives. To corroborate the *findsoloLTR* results, we interrogated the Database of Genomic Variants (DGV) (59) to assess whether any of the candidate dimorphic HERV-K or HERV-H loci had been previously predicted as copy number variants in the human population. The DGV systematically catalogs structural variants in human genomes reported in prior studies, but importantly it does not yet include data collected from the SDGP (55), thereby potentially serving as independent validation of our predictions from that dataset. We found that two of the four HERV-K candidates and more than half (35 out of 67) of the HERV-H candidates were catalogued in DGV as putative deletion variants (Table 1, Table S2). One of the HERV-K-associated deletions and twenty of the 35 HERV-H-associated deletions were inferred to have breakpoints mapping within the proviral LTRs, consistent with the idea that LTR recombination events caused these deletions (Table 1). The second HERV-K deletion reported in DGV has both breakpoints precisely at the outer boundaries of LTRs, which is consistent with a pre-integration allele previously reported (27). The remaining 15 HERV-H-associated deletions catalogued in DGV have predicted breakpoints mapping outside of the annotated LTR sequences, which suggests that a different mechanism than LTR recombination could have caused the deletion or that previous breakpoint identification might have been imprecise.

To further validate the *findsoloLTR* results, we selected one HERV-K candidate (5p13.3_K2) and two HERV-H candidates (2q34_H4, 3p14.3_H1) for experimental validation using PCR with primers designed in the flanking regions. In all three cases, the predicted solo LTR alleles were successfully detected by PCR and sequencing (Figure 2D-F), (Table 1, Table S2),(Supplementary Data1). Collectively these data demonstrate that the *findsoloLTR* pipeline efficiently predicts dimorphic HERVs and reveal that a surprisingly high fraction (up to ~10%) of HERV-H proviruses occur as solo LTR alleles in the human population, albeit at relatively low frequency.

### Potential consequences for transcriptome variation

To begin exploring the functional consequences of these structural variants, we sought to examine whether the candidate dimorphic HERVs were associated with any known protein-coding or non-coding genes (see methods). We found that three HERV-H candidates contribute exonic sequences including transcription start sites or polyadenylation signals to different RefSeq genes and 10 additional HERV-K and HERV-H loci contribute long intergenic non-coding RNA transcripts annotated in the human reference genome (Table S2). Furthermore, 52 of the HERV-H proviruses we predict to occur as solo LTRs in the population have been previously reported as either moderately or highly transcribed in human induced pluripotent stem cells (60). One of these HERV-H loci, which we validated experimentally (Figure 2F) corresponds to the RefSeq gene *Embryonic Stem cell Related Gene* (*ESRG*), which has been identified as a marker of pluripotency (60–63). The *ESRG* transcript initiates within the 5’ LTR of HERV-H and parts of its first and second exons are derived from the internal region of the element (60–62). Thus, it is likely that recombination to solo LTR would impair *ESRG* transcription and most likely its function. While preliminary, these observations suggest that HERV dimorphisms create structural variation that has the potential to impact the human transcriptome.

## DISCUSSION

Sustained efforts have been undertaken to map structural variation across human genomes in the general population or in association with diseases. But relatively sparse attention has been given to the identification of structural variants associated with HERVs, and particularly the type of dimorphism investigated in this study in which the ancestral allele is a provirus and the derived allele is a solo LTR. Such dimorphisms are challenging to identify because the two variants share the exact same junctions with flanking host DNA, which prevents their identification using ‘standard’ approaches based on split and discordant read mapping (e.g. (26, 52–54)). Here we have developed two pipelines that circumvent these challenges and efficiently identify dimorphic HERVs (Figure 1 D,E). Both pipelines rely on *a priori* knowledge of insertion sites in the reference genome and make use of paired-end and read depth information to infer whether a locus annotated as a provirus in the reference genome exist as a solo LTR in a sequenced individual and vice versa. Hence our approach differs from but complements previous efforts to identify HERV insertional polymorphisms (presence/absence), which by design cannot typically differentiate proviruses from solo LTRs (26, 52–54).

We applied our pipeline to discover dimorphic loci from three major HERV families of different ages (HERV-K, HERV-H, HERV-W) using sequence data generated from 279 individuals from diverse populations (55). Previously, only a dozen HERV-K insertions have been reported to exist as dimorphic provirus/solo LTR alleles in the human population (25–27, 35, 36, 40, 41, 43). Our results yielded 15 strong candidate HERV-K dimorphic loci, including 10 previously recognized as dimorphic in the human population, a subset of which are also known to be insertionally polymorphic (see Table 1, Table S1, Table S2) (25–27, 33–44). These results indicate that our approach is quite specific since it did not yield an extensive set of HERV-K candidates that were not identified previously. In turn, this observation suggests that the number of HERV-K loci with dimorphic alleles segregating with relatively high frequency in the human population is rather small and it appears that most of these loci have now been identified. Of course it is possible, and even likely, that many more dimorphic HERV-K loci segregate at low frequency in the population. While the SDGP represents a fairly diverse sampling of the human population compared to those previously surveyed for HERV polymorphisms such as the 1000 Genome Project, it still remains minuscule. As sequencing efforts continue to intensify worldwide, our pipeline brings a valuable addition to the toolbox for cataloguing structural variants.

We were intrigued to discover a dimorphic element for the HERV-W family (18q21.1_W2). This element is represented as a solo LTR in the reference genome, but our data clearly show that it also occurs as a provirus segregating in South Asian populations (Figure 2A) and likely in other diverse populations (our pipeline predicted a provirus allele in 194 out of 279 individuals surveyed, Table 1, Table S1). To the best of our knowledge, this is the first HERV-W locus reported to show any type of dimorphism. This particular HERV-W insertion must have occurred between 18 and 25 million years ago because a provirus is found at orthologous position in all other ape genomes including gibbon, but is absent in Old and New World monkeys (64). Our discovery illustrates the potential of LTR recombination to alter genome structure long after a proviral insertion has occurred.

We also identified an unexpectedly large number (~70) of candidate HERV-H dimorphisms. We experimentally validated the dimorphic nature of four of these HERV-H loci in South Asian populations and in an African individual (Table 1, Table S1, Table S2, Figure 2). While this is a small validation sample, the results suggest that a substantial number of HERV-H loci occur as dimorphic alleles in the human population, with solo LTR alleles apparently segregating at low frequency relative to proviral elements (Table 1, Table S1, Table S2). To our knowledge, prior to this study only a single dimorphic HERV-H locus had been documented (24). We did not identify this particular locus in our analysis. However, we noticed that the 5’ and 3’ LTRs of this provirus are annotated by Repeatmasker as belonging to different subfamilies (LTR7 and LTR7Y respectively), an annotation either erroneous or reflecting an inter-element recombination event (65). In either case, this discrepancy would have excluded this locus from our analysis because the program we used (66) to assemble the starting set of queries requires 5’ and 3’ LTR names to match in order for a locus to be flagged as a provirus. This observation highlights a caveat of our approach: it relies on accurate pre-annotations of the elements in a reference genome in order to correctly identify proviral and solo LTR queries. Clearly, repeat annotation remains an imperfect process even in a ‘reference’ genome, and HERVs and other LTR elements pose particular challenges for both technical and biological reasons (65, 67, 68). Efforts are underway to automate and improve repeat annotation (56, 69–72) as well as projects to enhance the quality of genome assemblies and annotations for a wide variety of species. These developments are bound to facilitate and expand the application of our pipeline to many more genomes, both human and non-human.

The unexpectedly large number of dimorphic HERV-H loci we predict in the population probably relate to some of the peculiar features of this family. HERV-H is a relatively abundant family that is exceptional for the high proportion of proviral insertions relative to solo LTRs maintained in the genome (73). By our estimates (see Methods) the reference genome includes ~720 HERV-H proviral insertions and 689 solo LTRs. Phylogenetic modeling of the LTR recombination process (73) suggests that HERV-H proviruses have formed solo LTRs at a much lower rate than expected based on their age of residence and the level of sequence divergence of their LTRs. Indeed HERV-K, a younger family, includes 23 proviral copies and 553 solo LTRs (see Methods). The apparent resistance of HERV-H to LTR recombination may be driven by purifying selection to retain proviral HERV-H copies for some sort of cellular function (73). In fact it has been documented that a subset of HERV-H proviruses are bound by pluripotency transcription factors and are highly expressed in human embryonic stem cells as long noncoding RNAs and chimeric transcripts playing a possible role in the maintenance of pluripotency (60, 74–77). Our finding that several HERV-H proviruses are reduced to solo LTR alleles in some individuals argues that haploidy for the internal sequences of these elements is sufficient for normal human development. But that is not to say that such structural variation bears no biological consequences. In fact, one of the dimorphic HERV-H loci we validated at 3p14.3 is known to drive *ESRG*, a transcript acting as an early marker of reprogramming of human cells to induced pluripotent stem cells (60–63). Experimental knockdown of the *ESRG* transcript in human embryonic stem cells leads to a loss of pluripotency and self-renewal (60). Thus, it is intriguing that we identified a solo LTR allele of *ESRG* in two individuals from different African populations (Table 1, Table S2, Figure 2F). Whether this deletion event impairs ESRG transcription and has any functional consequences for human embryonic development awaits further investigation. More generally, our catalog of candidate dimorphic HERVs provides a valuable resource to assess the regulatory significance of these type of elements (11) and assess whether the process of LTR recombination represents a hitherto ‘hidden’ source of regulatory divergence in the human population.

These findings also bear important implications for studies that link the coding activities of HERVs to human pathologies. Our results imply that there are more frequent alterations in the copy number of HERV coding sequences than previously appreciated, even for families that apparently have long ceased to be infectious or transpositionally active such as HERV-H and HERV-W (78, 79). Overexpression of gene products encoded by these families as well as HERV-K has been documented in a number of conditions, including multiple sclerosis (MS) (18), amyotrophic lateral sclerosis (ALS) (22), rheumatoid arthritis (80), systemic lupus erythematosus (81), schizophrenia (82) and type 1 diabetes (83) and several cancers (84–87). It remains uncertain whether overexpression of HERVs contributes to the etiology or progression of these diseases. But evidence is mounting in the cases of MS and ALS, for which both *in vitro* studies and mouse models have established that envelope (*env*) proteins expressed by HERV-W and HERV-K respectively, can exert biochemical, cellular and immunological effects that recapitulate the disease symptoms (18). Conceivably then, variation in the copy number of HERV-encoded genes caused by sporadic LTR recombination events, either in the germline or in somatic cells, could modulate susceptibility to these pathologies. Importantly, three of the dimorphic HERV-K loci predicted herein (Table1, Table S2) are known to encode full-length env proteins (88). Thus, our results reveal a previously underappreciated source of HERV gene copy number variation with potential pathological ramifications.

Lastly, a growing number of studies have implicated HERV-encoded proteins in beneficial physiological activities, notably in immunity (for review (10)). For instance, overexpression of the HERV-K *gag* protein can interfere with the late phase replication of the HIV-1 retrovirus (89). Moreover, biochemically active HERV-K proteins appear to be expressed during normal human development where they may confer some form of immunity to the early embryo (90, 91). For example, endogenous *env* can compete with and effectively restrict the cellular entry of cognate exogenous retroviruses (92, 93), and *env* of the HERV-H and HERV-W families have been shown to have immunosuppressive properties (94, 95). Thus, it is tempting to speculate that some of the genomic variants uncovered herein could contribute to inter-individual immune variation and modulate the risk to develop certain pathologies.

## CONCLUSIONS

Collectively our results show that we have successfully developed a pipeline to discover dimorphic loci from a variety of HERV families from resequencing data, including two families for which such copy number variation had been scarcely (HERV-H) or never (HERV-W) reported before. Given that there are dozens more HERV families in the human genome, including some substantially younger than HERV-H or HERV-W (65, 68), it is likely that this form of structural variation affect other families and is more common than previously appreciated. Further studies are warranted to investigate the association of such variants with human phenotypes, including disease susceptibility.

## METHODS

### Classification of proviruses and solo LTRs in the reference genome

The repeats annotated as LTR5-Hs and HERV-K-int (HERV-K(HML2 family)), as LTR17 and HERV17-int (HERV-W family) and as LTR7 and HERV-H-int (HERV-H family) are extracted from the RepeatMasker annotation of the human reference (GRCh38/hg38) assembly (RepeatMasker open-4.0.5 - Repeat Library 20140131 available at http://www.repeatmasker.org/). The extracted RepeatMasker data is parsed to identify potentially full-length proviruses and solo LTRs using the tool “One Code to Find Them All” (66). Using a custom script, (https://github.com/iainy/dimorphicERV) each copy in the parsed output is further classified as a provirus containing (i) 2 LTRs and internal region (ii) 1 LTR and internal region (iii) only internal region or as a solo LTR. The coordinates at the boundaries of each copy is then extracted from the parsed output. Each HERV locus is then given a unique identifier depending on the cytoband it belonged to and based on the total number of copies of that family found in each band. The positions of cytoband for GRCh38/hg38 is downloaded (http://hgdownload.cse.ucsc.edu/goldenpath/hg38/database/cytoBand.txt.gz). The coordinates of HERV copies marked as proviruses with 2LTRs and internal regions and as solo LTRs are used in the subsequent analysis. For HERV-W, the copies that are generated by retrotransposition mediated by LINE-1 machinery have partial LTRs (96) and such copies annotated as pseudogenes (78) were excluded from our analysis.

### Identification of provirus allele when the reference allele is a solo LTR

The *findprovirus* pipeline identifies solo LTR to provirus variants in the Binary Alignment/Map (bam) format files where paired end reads from whole genome resequencing data are mapped to reference assembly using Burrows-Wheeler Aligner (BWA) (97) (Figure 1D, Figure S1) (https://github.com/jainy/dimorphicERV). The pipeline analyses the coordinates of all solo LTRs obtained from One Code to Find Them All (see methods). The *findprovirus* pipeline extracts reads mapped to each solo LTR and to a flanking 100-bp region using samtools (version 1.4.1) (98). Only reads that are mapped with a mapping quality of 30 or greater (i.e. mapped with >99.99% probability) are collected and the reads are processed to fasta format using SeqKit (99). The discordant reads in the solo LTR and in the flanking 100-bp region are identified using samtools (98) and the mates of discordant reads are extracted using picard tools (version 2.9.2) (http://broadinstitute.github.io/picard/). Sequence homology of mates of discordant reads to the consensus coding sequence of the respective HERV extracted from the Repbase database (56) is tested using BLASTn (version 2.6.0, default parameters) and the number of reads with significant hits (e-value < 0.0001) are counted. Discordant read mates with significant sequence similarity to an internal HERV sequence suggest the presence of a proviral allele in that individual. Two independent, additional approaches are also used to assess the presence of a proviral allele. The first approach measures the average read depth at the solo LTR using samtools and reads with a mapping quality of 20 or more (mapped with > 99% probability) and reads with a base quality of 20 or more (base call accuracy of > 99%) are counted. To get an estimate of the expected coverage at a solo LTR, average of read depths at all solo LTRs of that HERV family for an individual is calculated. This also helps to account for the variability in the coverage between individual genomes. The ratio of average read depth at a solo LTR to the average of read depths observed at all solo LTRs of that HERV family for the individual is determined. An increased read depth pertained to the solo LTR (ratio > 1) is indicative of an increased number of reads mapping to that locus, which is suggestive of the presence of a provirus allele (Figure S1). As part of the second approach, a local *de novo* assembly of all extracted reads from a locus (mapped reads and discordant mates) is performed using CAP3 (100) and/or SPAdes (version 3.11.1) (101) to test if the solo LTR allele could be reconstructed. The corresponding reference solo LTR sequence with 50-bp flanking is extracted and sequence similarity of the reference sequence is tested (BLASTn version 2.6.0, default parameters) against assembled contigs. A significant blast hit (e-value < 0.0001) spanning ⩾95% reference genome sequence is indicative of the presence of a solo LTR allele in the individual examined. In addition to these two approaches, to discern a strong candidate from weak candidate, the mappability of genomic regions (102) where informative discordant reads are mapped is determined for each locus. The regions of low mappability generate ambiguous mapping and regions of high mappability generate unique mapping. The mappability scores are downloaded for the GRCh37/hg19 version of reference assembly (ftp://hgdownload.soe.ucsc.edu/gbdb/hg19/bbi/wgEncodeCrgMapabilityAlign100mer.bw). The downloaded file is processed (103) and is converted to bed format (104) and scores are lifted over (105) to hg38 version. This data is stored in an indexed mysql table. The coordinates of the reference assembly where the informative discordant reads are mapped for each solo LTR are identified using bedtools (version 2.26.0) (106). The mappability scores for those genomic regions are extracted from the table and the mean of the mappability scores is provided in the output of the pipeline.

### Identification of solo LTR allele when the reference allele is a provirus

The *findsoloLTR* pipeline identifies the provirus to solo LTR variants in bam files (Figure 1E, Figure S3) (https://github.com/jainy/dimorphicERV). It first calculates the read depth across the provirus using samtools (98). Read depth is calculated for reads with a mapping quality of 30 or more and with a base quality score of 20 or more. Similarly, read depth is calculated across 5’ and 3’ flanking 250-bp regions. The pipeline then assess the percentage of average read depth across the provirus to the average of read depths across the flanks. Presence of two proviral alleles is inferred when the read depth percentage greater than or equal to 50% and read depth percentage lower than 50% is used to infer the presence of solo LTR allele (Figure 1E). A read depth percentage lower than 10% is arbitrarily used to infer the presence of two solo LTR alleles. The mappability scores (102) of the genomic region spanning the the provirus are extracted (see methods for *findprovirus*) and the mean of the mappability scores is provided in the output of the pipeline.

### Dataset analyzed

The two pipelines were ran on the publicly available whole genome sequence data generated as part of the SGDP for 279 individuals from 130 populations (55). The bam files generated by aligning 100-bp long paired-end reads to the GRch38/hg38 version of the human genome using BWA aligner (version 0.7.12) (97) are used for the analysis.

### *In silico* validation

An *in silico* validation of the candidates identified by both pipelines is performed to filter out false positives. Each of the candidate loci including their flanking region (1000 bp) was visually inspected using IGV (version 2.3.97) after loading a track with RepeatMasker annotation of hg38 version of the human genome (RepeatMasker open-4.0.5 - Repeat Library 20140131). The candidates (identified through *findprovirus* pipeline) having an internal region of the respective HERV family nearby or having a nested polymorphic TE, both hallmarks of false positives, are filtered out. Candidate loci not supported by a minimum of four discordant reads where mates align to the internal coding sequence of HERV in at least one individual are also filtered out. The candidates (identified through *findsoloLTR* pipeline) having deletion restricted to a fragment of internal sequence are removed. After visual inspection, the candidates are then queried in the DGV (59) to identify if any previous studies have reported those loci as a copy number variant (CNV). The CNVs identified in DGV are visually inspected for the concordance of their breakpoints with the two LTRs, which is suggestive of their origin through a LTR mediated recombination. The CNVs having one or both breakpoints lie outside the LTRs are also identified. The candidates along with 100-bp flanking sequence are also queried against nr/nt database at NCBI to identify the presence of any BAC/FOSMID clones containing corresponding the solo LTR or provirus variant.

### Experimental validation

After *in silico* validation, PCR primers are designed in the regions flanking the LTR and in the *gag* and/or *env* regions assembled from the mates of the discordant reads for selected candidates. The solo LTR allele is amplified by primer pairs flanking the solo LTR and the proviral allele is amplified with the internal primer located on the *env* region or *gag* region. The primers for validating the dimorphic HERVs are designed using PrimerQuest (107) and the oligos are synthesized from Integrated DNA Technologies (IDT). For PCR validation, genomic DNA samples are selected based on the predicted genotype and availability. The sample ids of 12 individuals in the SGDP data set (55) used for PCR analysis are S_Brahmin-1, S_Brahmin-2, S_Irula-2, S_Kapu-1, S_Kapu-2, S_Madiga-2, S_Mala-2, S_Mala-3, S_Relli-1, S_Yadava-1, S_Yadava-2 and S_Luhya-2. PCR amplifications are performed using GoTaq PCR Master Mix (Promega) or Platinum SuperFi PCR Master Mix (Thermo Fisher Scientific). The primer sequences and PCR conditions used for each reaction are given in Table S3. PCR products are visualised using agarose gel electrophoresis and are purified using DNA Clean & Concentrator™-5 (Zymo Research) following manufacturer’s instructions. The purified PCR products are Sanger sequenced at the DNA sequencing Core Facility, University of Utah or at Genewiz. The generated sequences are analysed using Sequencher 5.4.6 (Gene Codes Corporation).

### Analysis of contribution of dimorphic candidate HERVs to annotated genes/transcripts

The dimorphic candidate HERV loci are examined individually using the University of California, Santa Cruz (UCSC) genome browser on human GRCh38/hg38 assembly (108) (last accessed June 6 2018) to identify any overlap with known NCBI RefSeq protein-coding or non-coding genes (NM_*, NR_*, and YP_*). In addition, to determine the dimorphic candidates that encode an intact *env* gene, the HERV coordinates are compared with that of intact *env* Open Reading Frames (ORFs) identified by Heidmann *et al*. (88) in the human genome (hg38). In order to find the candidate dimorphic HERV-Hs that are actively transcribed in human embryonic or induced pluripotent stem cells (iPSCs), coordinates HERV-Hs, which are known to be moderately or highly expressed in hiPSC lines and single cells (60) are intersected with coordinates of dimorphic HERV candidates using bedtools v2.26.0 (106).

## Acknowledgments

This work was supported by a research agreement between GeNeuro Innovation SAS and the University of Utah as well as funds from the National Institutes of Health to C.F. We would like to thank W. Scott Watkins for providing the 279 SGDP bam files for the analysis and Lynn B. Jorde for providing the DNA samples as well as equipment and infrastructure for performing this study. We also would like to thank Aurelie Kapusta and Clement Goubert for many helpful discussions.

## Supplementary Information

**Figure S1.**
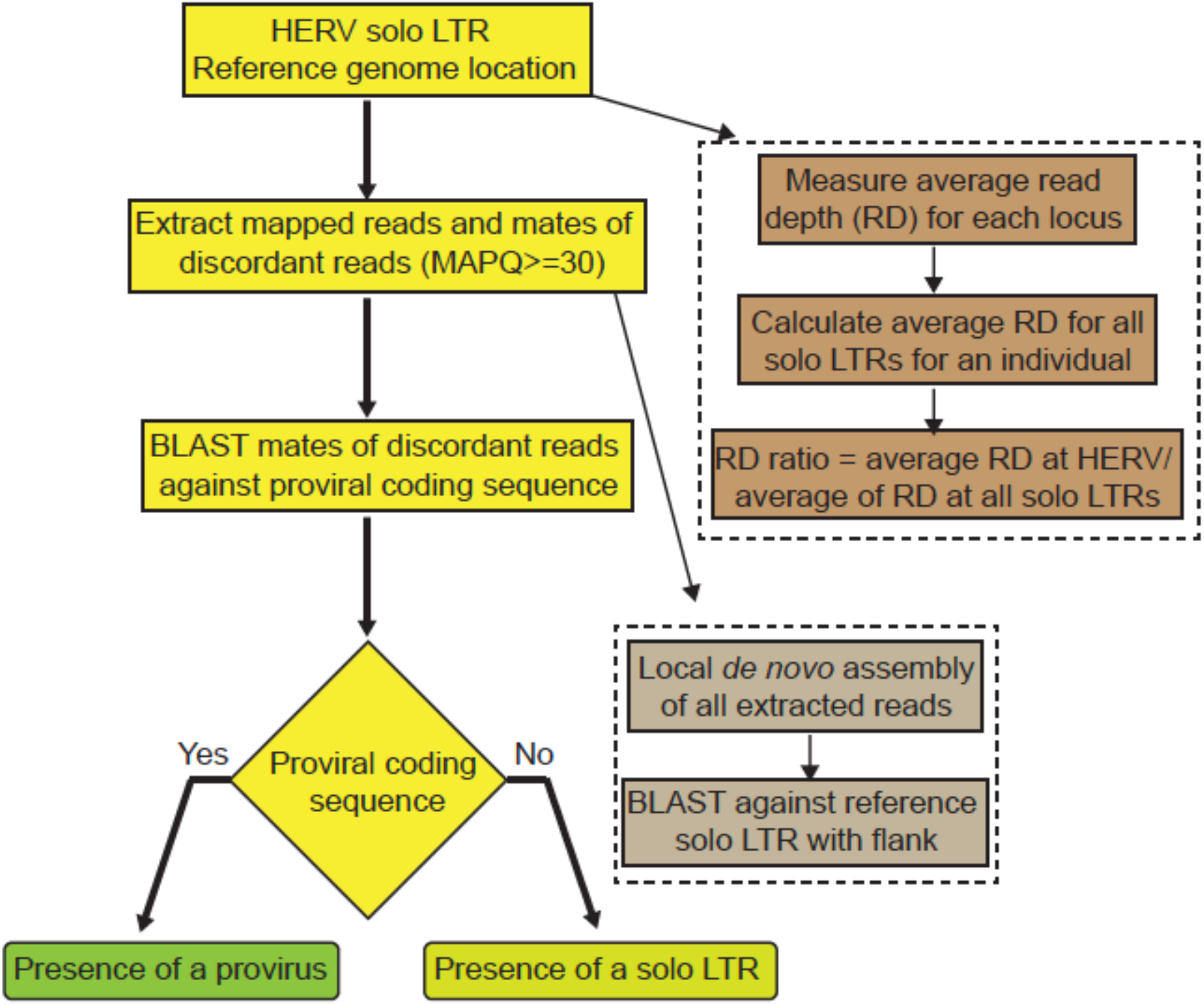
Flowchart of *findprovirus* pipeline. The first step indexes the coordinates of solo LTRs of a HERV family in the reference genome. Mapped reads (of mapping quality score (MAPQ) equal or greater than 30) and mates of discordant reads are extracted in a window extending ±100-bp from each LTR. Homology based searches are performed with mates of discordant reads against the respective consensus of internal sequence of HERV to infer the presence of a provirus allele at the locus. The read depth for each locus is calculated and compared to the average of read depths for all solo LTRs of that family in an individual. Increased read depth may be observed for some candidate loci reflecting the presence of a provirus allele. A local *de novo* assembly of the reads is also performed to infer the presence or absence of a solo LTR allele at the locus. These two additional approaches (enclosed by dashed lines) are performed by the pipeline but are not primarily used to infer the presence of a provirus.

**Figure S2.**
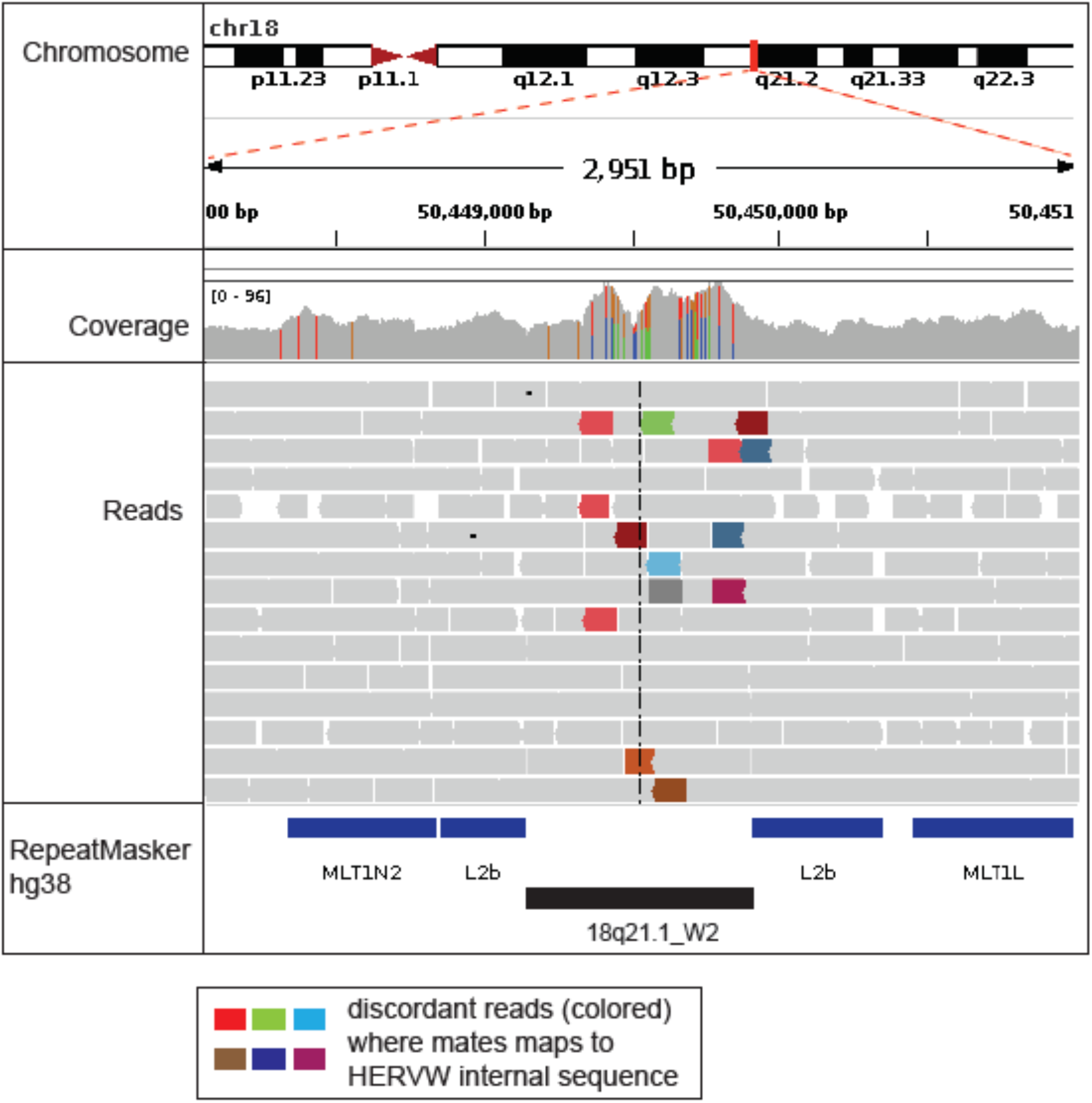
IGV screenshot of the dimorphic HERV-W locus 18q21.1_W2. The top panel shows the position on the chromosome and hg38 coordinates. The Coverage track shows the number of reads that covers each base at the locus. The numbers within brackets [0-96] shows the range of reads covering the visualized region. The Reads track shows a sample of the reads mapped to that locus. The colored reads are discordant reads whose mates have unexpected insert size or inconsistent orientation. The RepeatMasker track shows the position of transposable elements and repeats identified in that chromosomal interval. The presence of discordant reads with mates mapping to internal HERV-W coding sequence is used to infer the presence of provirus at the locus. The increase in read coverage pertaining to the solo LTR region is also indicative of presence of provirus, as the reads from 2 LTRs of a provirus allele maps to the solo LTR. The candidate is experimentally verified to be homozygous for provirus at that locus.

**Figure S3.**
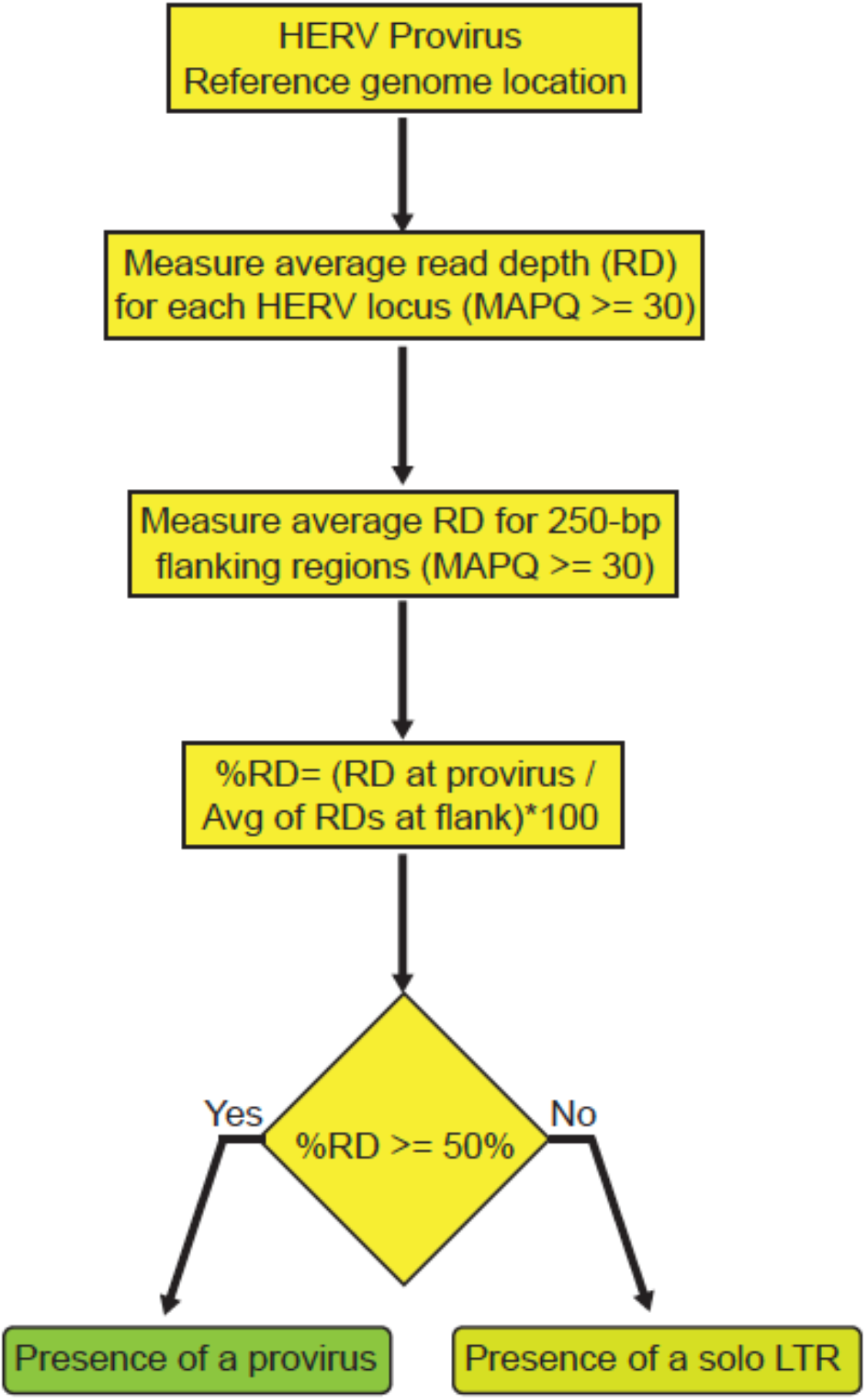
Flowchart of *findsoloLTR* pipeline. The first step indexes the coordinates of proviruses of a HERV family in the reference genome. Average of read depth (of mapping quality score (MAPQ) equal or greater than 30 and base call accuracy equal to or greater than 20) at the HERV locus and at the flanking window extending ±250-bp from both LTRs are calculated. Percentage of the average read depth at each HERV locus to the average of the read depths at the two flanking 250-bp window is assessed. An estimated percentage equal to or greater than 50% is used to infer the presence of a provirus and the percentage lower than 50% infer the presence of a solo LTR allele.

**Figure S4.**
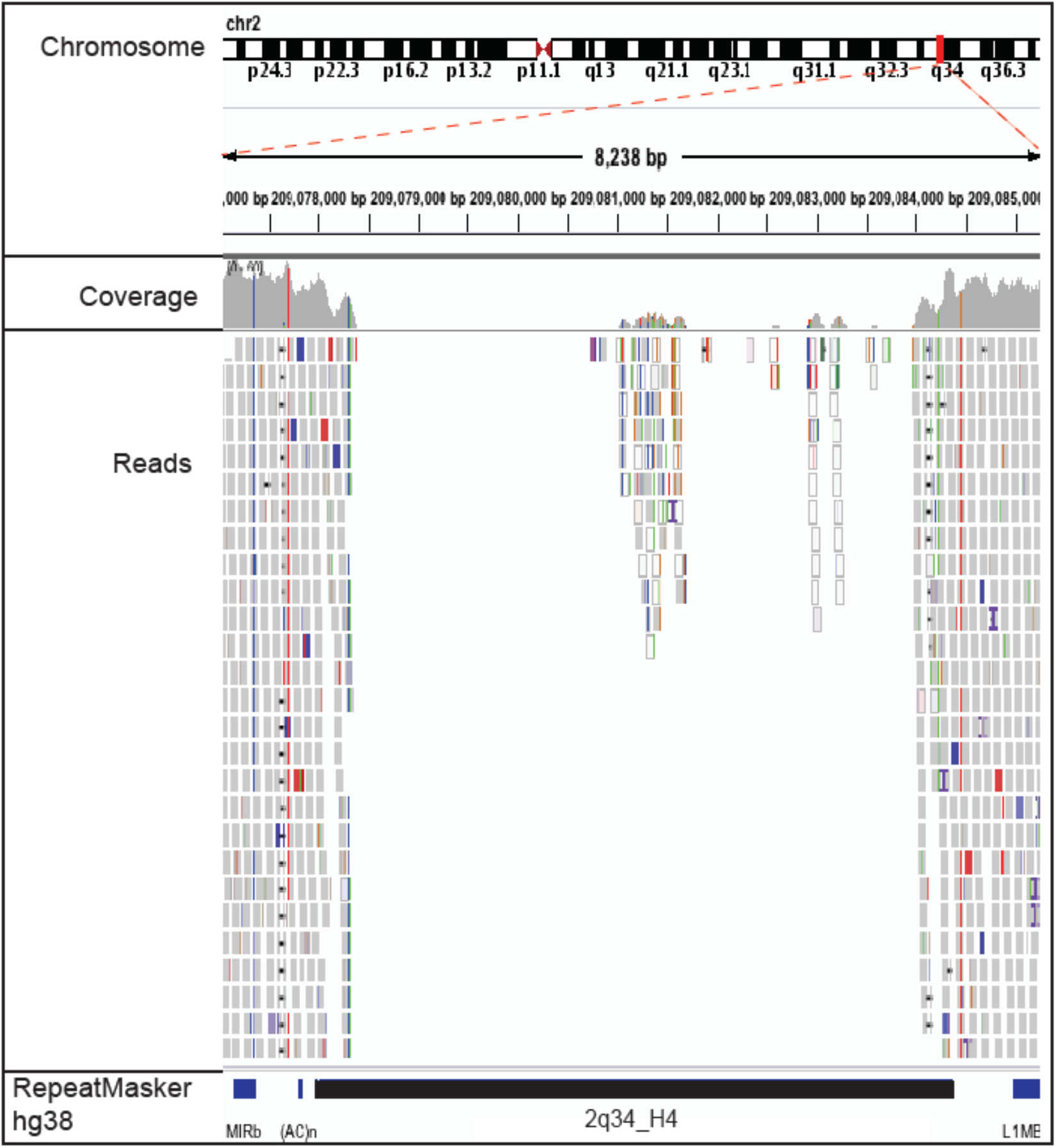
IGV screenshot of dimorphic HERV-H (2q34_H4) locus. The top panel shows the position on the chromosome and hg38 coordinates. The Coverage track shows the number of reads that covers each base at the locus. The numbers within brackets [0-60] shows the range of reads covering the visualized region. The Reads track shows a sample of the reads mapped to that locus. The RepeatMasker track shows the position of transposable elements and repeats identified in that chromosomal interval. Here, nearly a complete lack of uniquely mapping reads to the internal region suggest the presence of two solo LTR alleles.

**Table S1.**
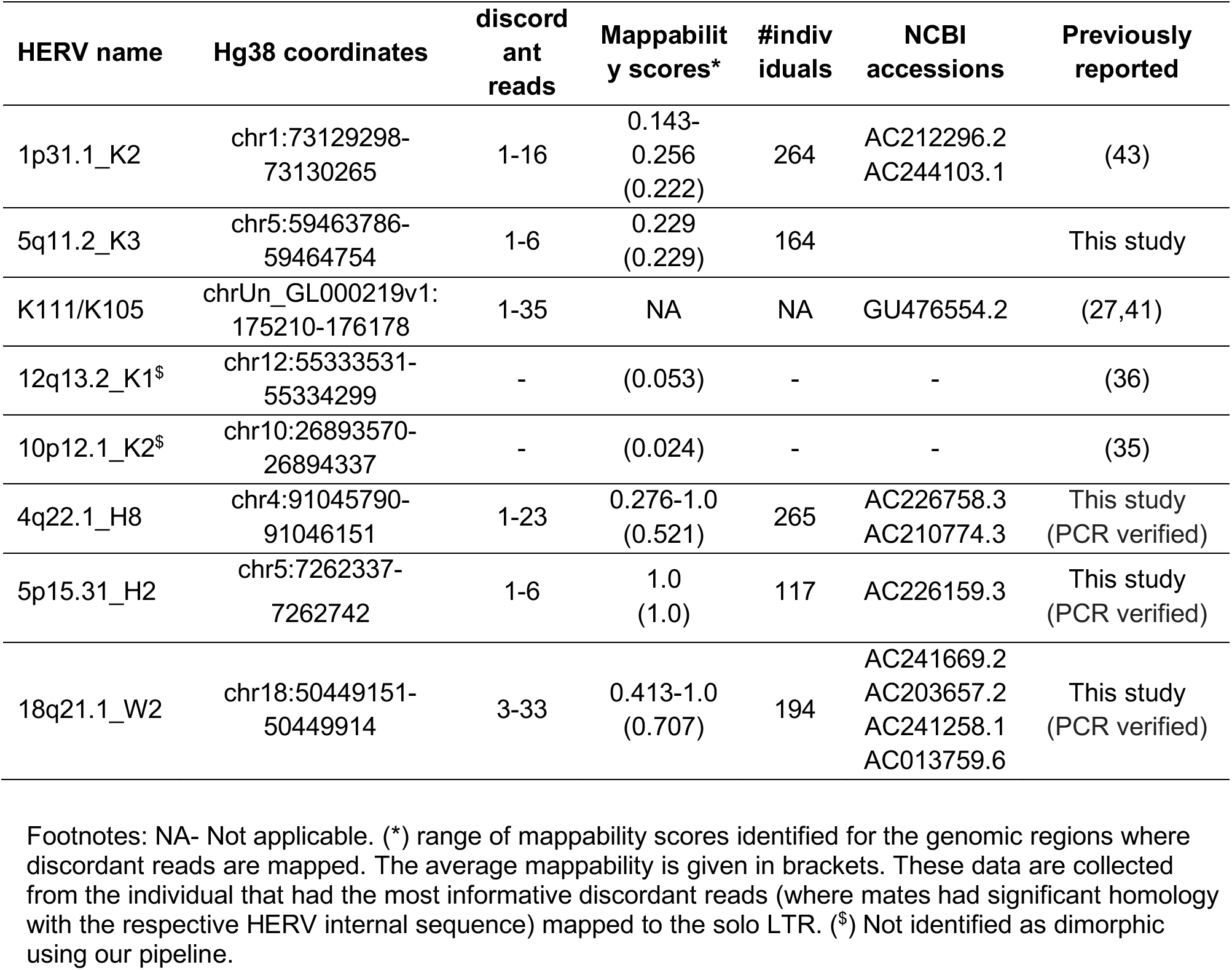
HERV candidates identified as solo LTR to provirus variants using *findprovirus* pipeline

**Table S2.**
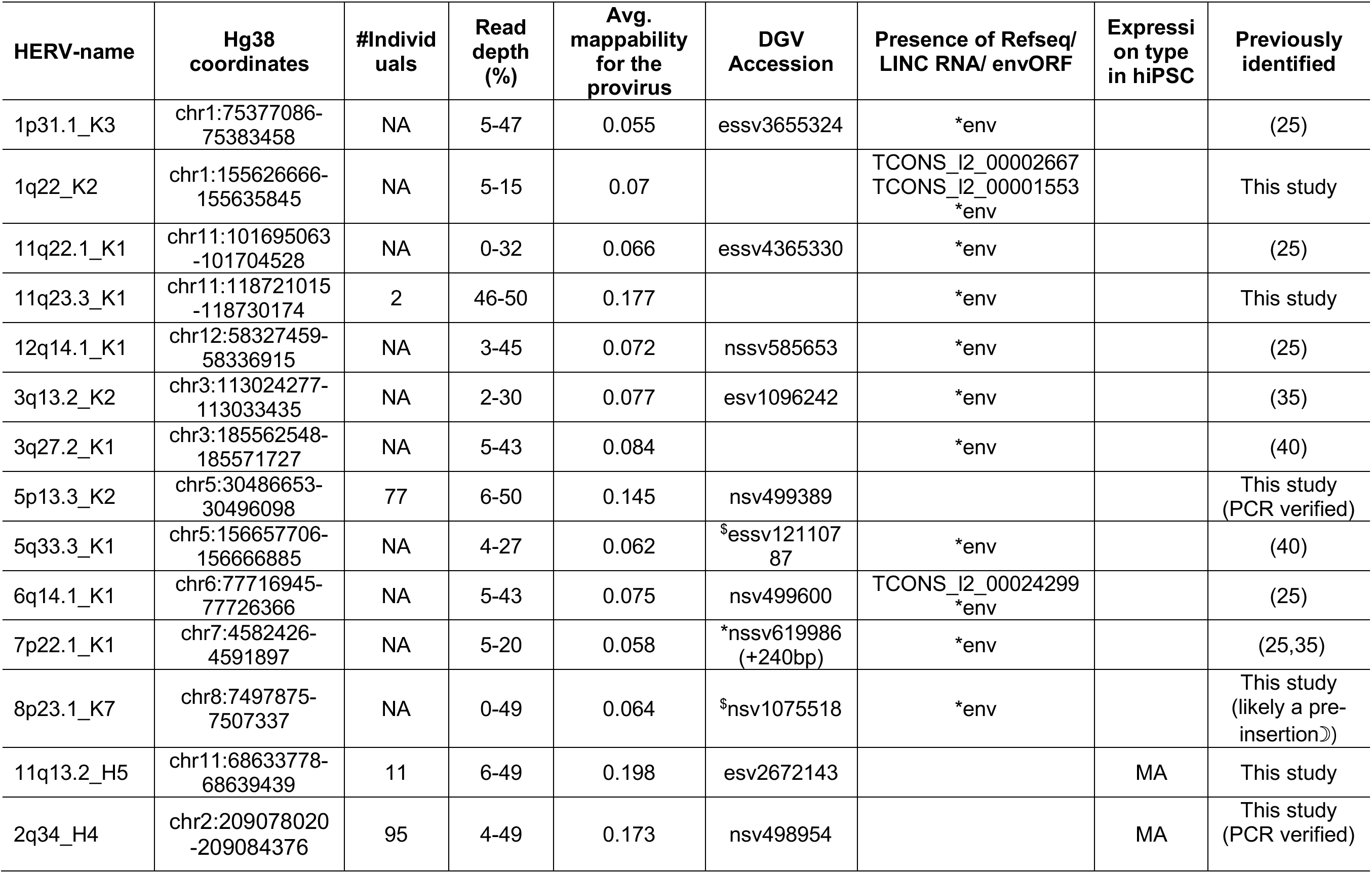
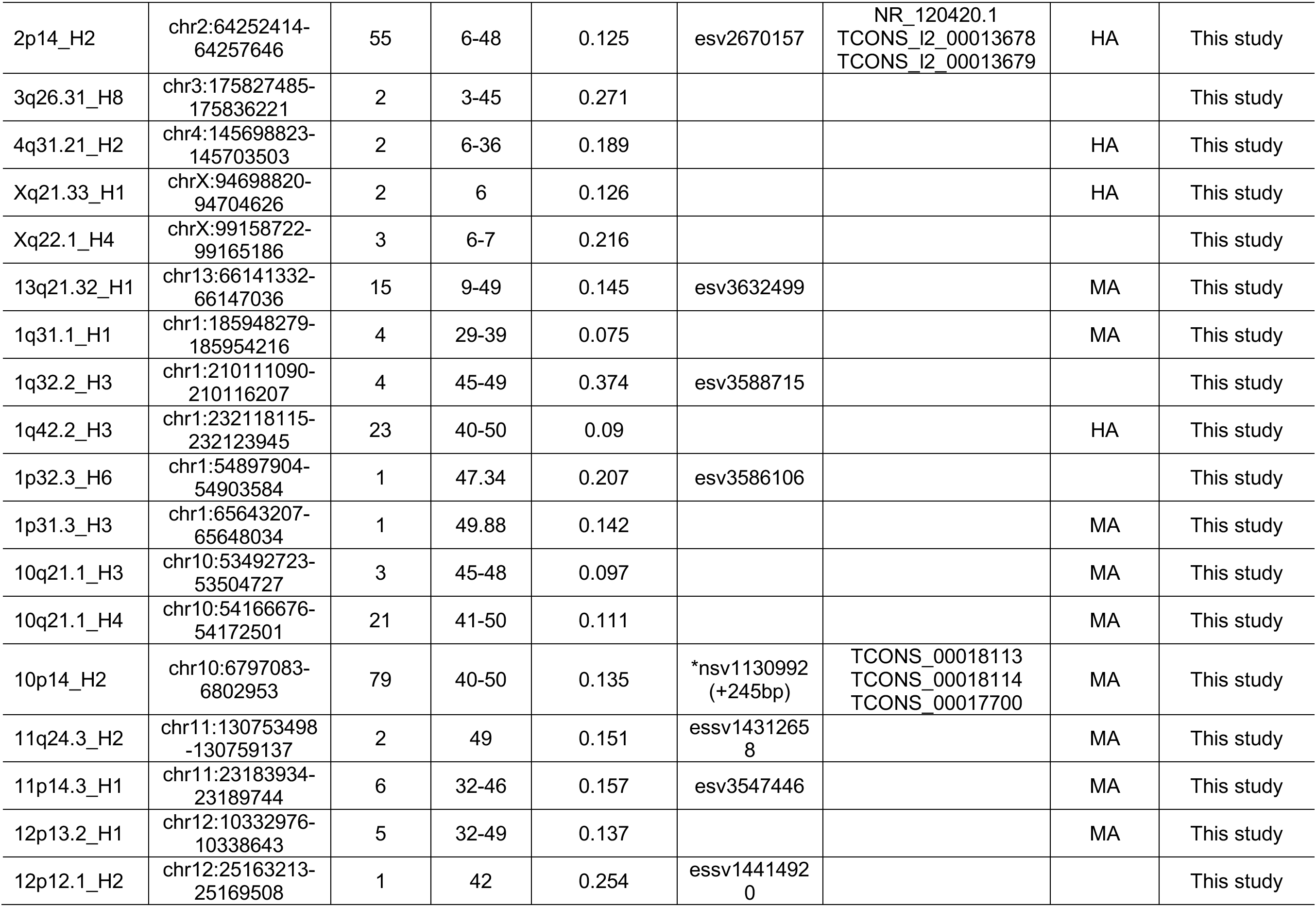
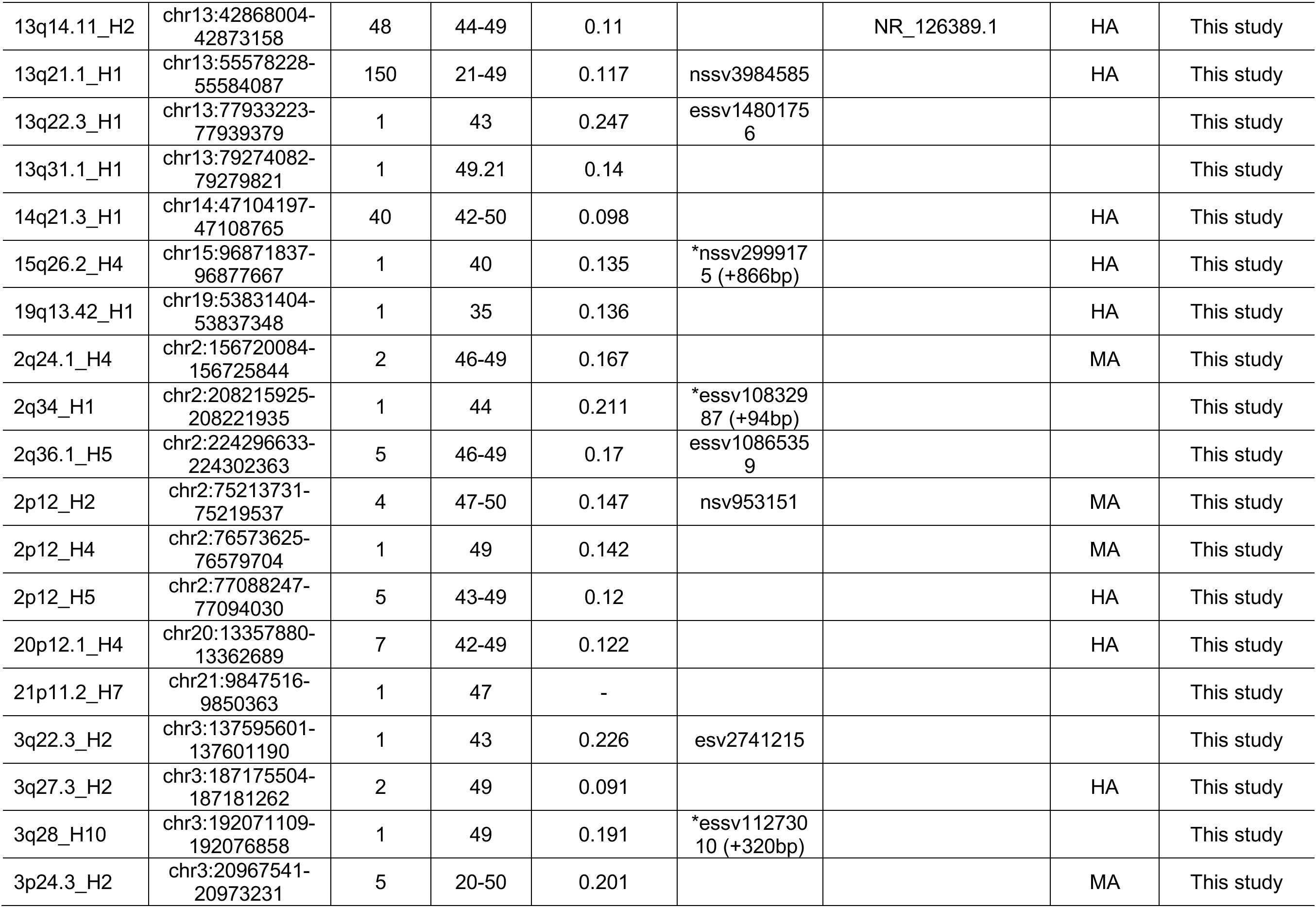
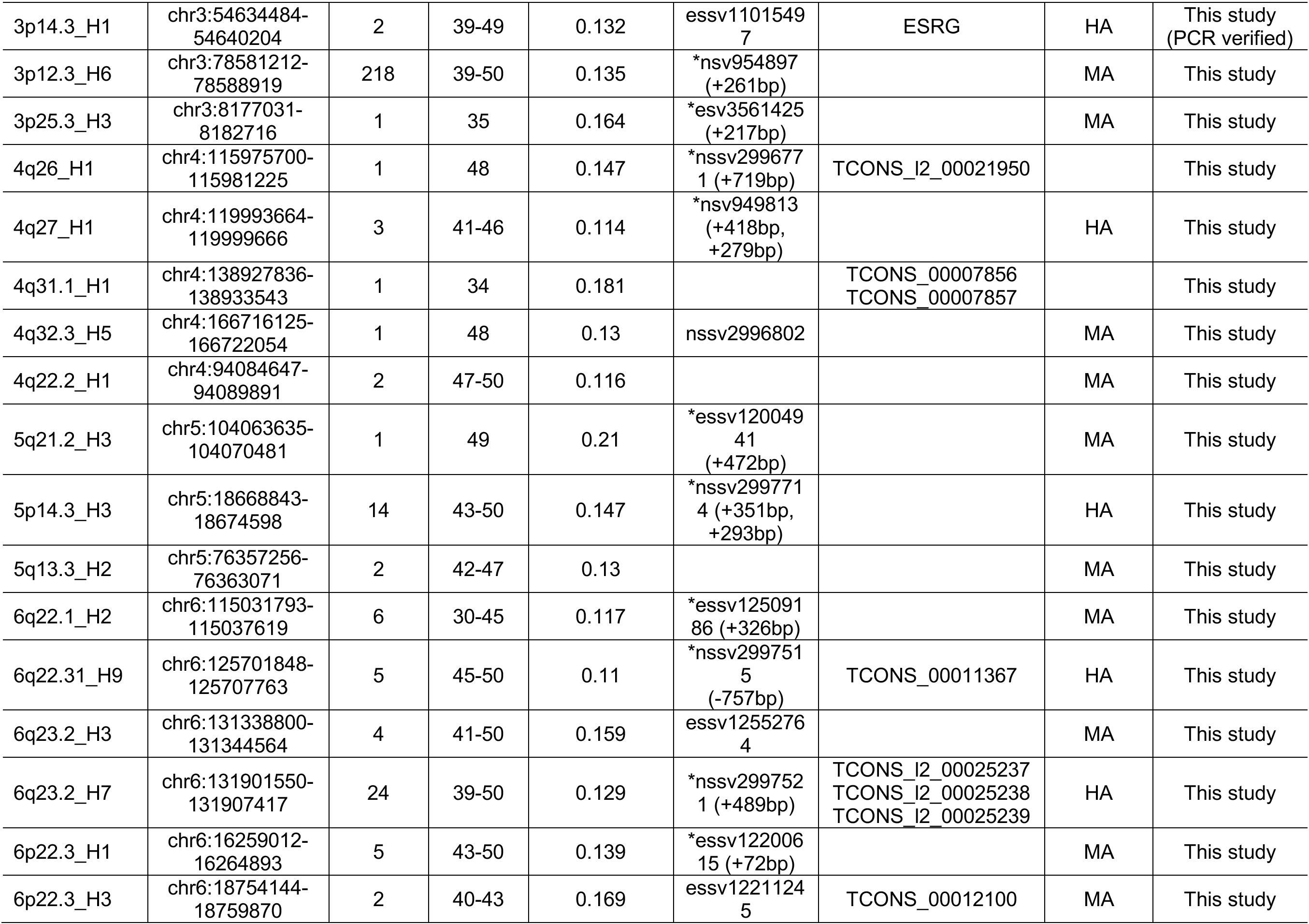
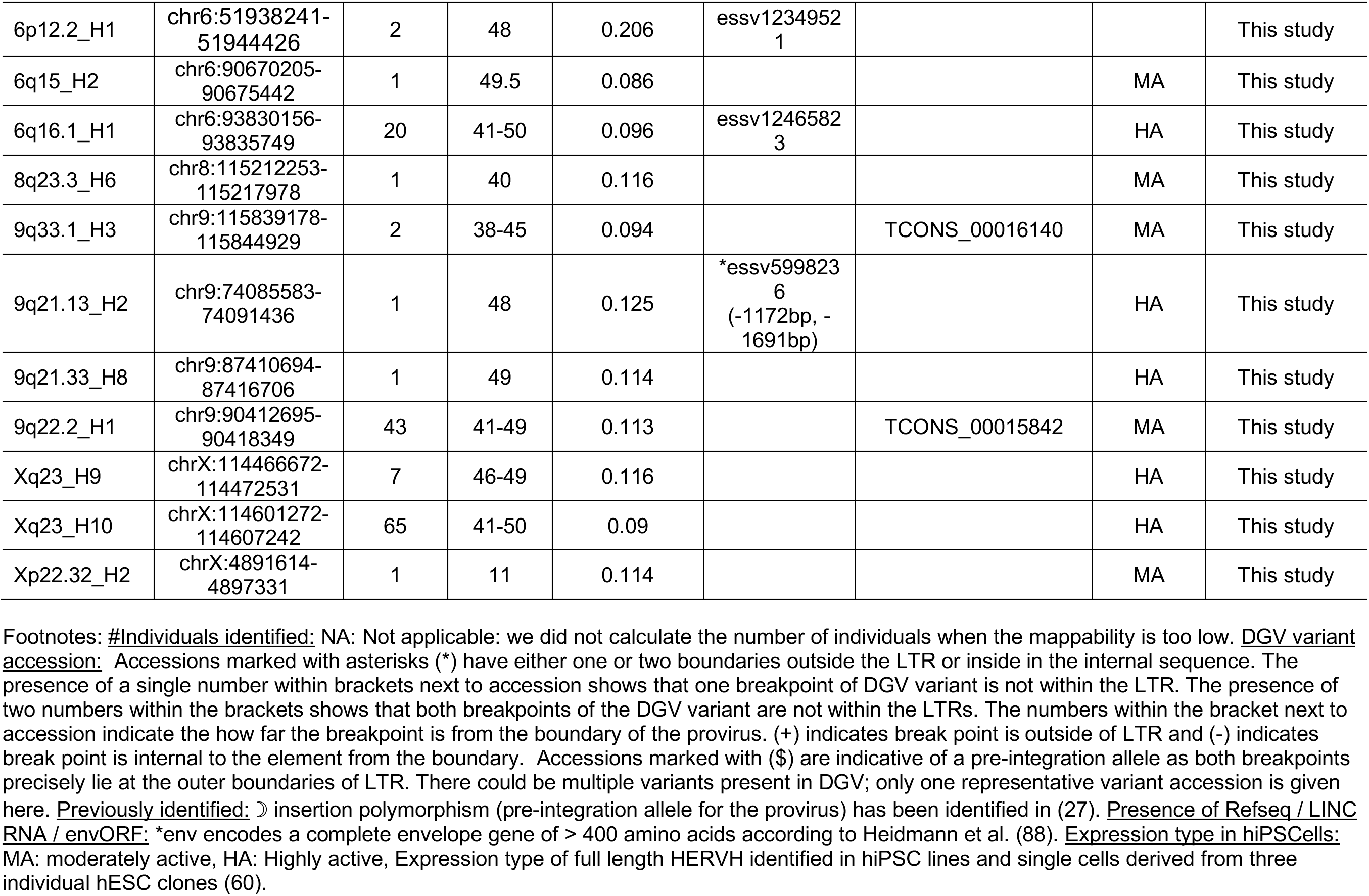
HERV candidates identified as provirus to solo LTR variants using *findsoloLTR* pipeline

**Table S3.**
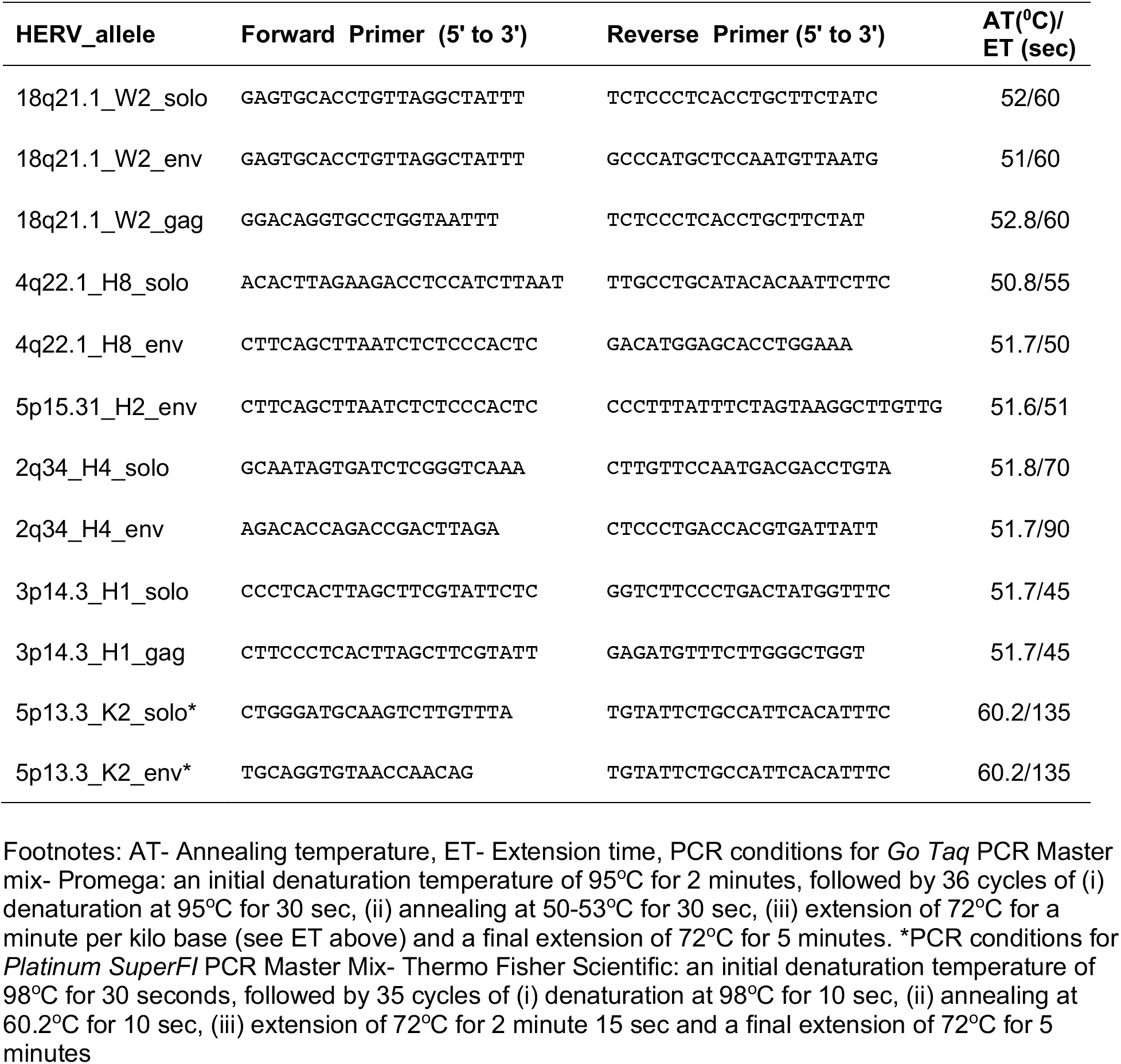
List of primer sequences used for amplifying solo LTR and provirus alleles shown in Figure 2

### Supplementary Data 1

`>S Yadava-1 18q21.1 W2 env partial sequence`

GGAAATCTCNCTGCACAACCCCTACTACACCCCAATTCAGCAGGAAGCAGTTAGAGCAGTTGTCAGCCAACCTCCCCAACAGCACTTGGGTTTTCCTGTTGAGAGGGGGGACTGAGAGACAGGACTAGCTGGATTTCCTAGGCCAACTAAGAATCCCTAAGCCTAGCTGGGAAGGTGACCGCATCCACCTTTAAACATGGGGCTTGCAACTTAGCTCACACCGGCCAACCAGGTAATAAAGAGAGCTCACTAAAATGCTAATCAGGCAAAAACAGCAGGTAAAAAAATAGCCAATCATCTTTCGCCTGAGACCACAGTGGGCGGGACAATGATCAGGATATAAACCCAGGCATTCAAGCCAGCAATGGCTACCCTCTTTGGGTCCCCTCCCTTTGTATGGGAGCTCTGTTTTCGCTCTATTAAATCTTGCAACTGCGCTCTCTTCTGGTGCATGTTTGCTATGGCTCGGCTCTAGCTGAGCTTTTGTTCACTGTCCACCACTGCTGTTTGCTGCCGCCGCAGACCTGCCACTGACTTCCATCCCTTCGGATTCGGCAGGGTGTCCTCTGTGCCCCTGATCCAGCAAGGCACCCATTGCTGCTCCTGATCAGGCTAAAGTCTTGCCATTGTTCCTGCATGGCTAAGTGCCCAGGTTCATCCTAATTGAGCTGAACACTAGTCACTGGGTTCACAGTTCTCTTCCGTGACCCATGGCTTCTAATAGAGCTATAACACTCACTGCATGGCCCAAGATTCCATTCCTTGGAATCCATGAGGCCAAGAACCCCNGTCAGAGAACACGAGGCTTGCCACCATTTTGGAAGTGGCCTGCCGCCATCTTGGGAGCTCTGGGANCAAGGACTCCCCCGGTAACACTACCATCATCTCTCATCTGATAAGTGAAATAGCCTA

`>S Yadava-2 18q21.1 W2 env partial sequence`

AATCTCNNTGCACAACCCCTACTACACCCCAATTCAGCAGGAAGCAGTTAGAGCAGTTGTCAGCCAACCTCCCCAACAGCACTTGGGTTTTCCTGTTGAGAGGGGGGACTGAGAGACAGGACTAGCTGGATTTCCTAGGCCAACTAAGAATCCCTAAGCCTAGCTGGGAAGGTGACCGCATCCACCTTTAAACATGGGGCTTGCAACTTAGCTCACACCGGCCAACCAGGTAATAAAGAGAGCTCACTAAAATGCTAATCAGGCAAAAACAGCAGGTAAAAAAATAGCCAATCATCTTTCGCCTGAGACCACAGTGGGCGGGACAATGATCAGGATATAAACCCAGGCATTCAAGCCAGCAATGGCTACCCTCTTTGGGTCCCCTCCCTTTGTATGGGAGCTCTGTTTTCGCTCTATTAAATCTTGCAACTGCGCTCTCTTCTGGTGCATGTTTGCTATGGCTCGGCTCTAGCTGAGCTTTTGTTCACTGTCCACCACTGCTGTTTGCTGCCGCCGCAGACCTGCCACTGACTTCCATCCCTTCGGATTCGGCAGGGTGTCCTCTGTGCCCCTGATCCAGCAAGGCACCCATTGCTGCTCCTGATCAGGCTAAAGTCTTGCCATTGTTCCTGCATGGCTAAGTGCCCAGGTTCATCCTAATTGAGCTGAACACTAGTCACTGGGTTCACAGTTCTCTTCCGTGACCCATGGCTTCTAATAGAGCTATAACACTCACTGCATGGCCCAAGATTCCATTCCTTGGAATCCATGAGGCCAAGAACCCCAGGTCAGAGAACACGAGGCTTGCCACCATTTTGGAAGTGGCCTGCCGCCATCTTGGGAGCTCTGGGAGCAAGGACTCCCCCGGTAACACTACCATCATCTCTCATCTGAATAAGTGAAATAGCCTAC

`>S Mala-3 18q21.1 W2 env partial sequence`

GGTAGTGTTACCGGGGGAGTCCTTGCTCCCAGAGCTCCCAAGATGGCGGCAGGCCACTTCCAAAATGGTGGCAAGCCTCGTGTTCTCTGACCTGGGGTTCTTGGCCTCATGGATTCCAAGGAATGGAATCTTGGGCCATGCAGTGAGTGTTATAGCTCTATTAGAAGCCATGGGTCACGGAAGAGAACTGTGAACCCAGTGACTAGTGTTCAGCTCAATTAGGATGAACCTGGGCACTTAGCCATGCAGGAACAATGGCAAGACTTTAGCCTGATCAGGAGCAGCAATGGGTGCCTTGCTGGATCAGGGGCACAGAGGACACCCTGCCGAATCCGAAGGGATGGAAGTCAGTGGCAGGTCTGCGGCGGCAGCAAACAGCAGTGGTGGACAGTGAACAAAAGCTCAGCTAGAGCCGAGCCATAGCAAACATGCACCAGAAGAGAGCGCAGTTGCAAGATTTAATAGAGCGAAAACAGAGCTCCCATACAAAGGGAGGGGACCCAAAGAGGGTAGCCATTGCTGGCTTGAATGCCTGGGTTTATATCCTGATCATTGTCCCGCCCACTGTGGTCTCAGGCGAAAGATGATTGGCTATTTTTTTACCTGCTGTTTTTGCCTGATTAGCATTTTAGTGAGCTCTCTTTATTACCTGGTTGGCCGGTGTGAGCTAAGTTGCAAGCCCCATGTTTAAAGGTGGATGCGGTCACCTTCCCAGCTAGGCTTANGGATTCTTAGTTGGCCTANGAAATCCAGCTAGTCCTGTCTCTCAGTCCCCCCTCTCAACAGGAAAACCCAAGTGCTGTTGGGGAGGTTGGCTGACAACTGCTCTAACTGCTTCCTGCTGAATTGGGGTGTAGTAGGGGTTGTGCAGTTGAGATTTCCTTGGGAGGGTGCNTTGATGTCAT

`>S Kapu-1 18q21.1 W2 gag partial sequence`

GANAGCAAGCGNAAGAGGTCCAATATTACTCACTGCTTTGGAGATCTCTTCGTGGTTACCAAAATGTCACCGGGGGTTCCTTGCTCCCAGAGCTCCCAAGATGGCAGCAGGCCACTTCCAAGATGGTGGCAAGCCTTGGCCTCACGGATTCCAAAGAATGGAATCTTGCACCATGCGATGAGTGTTATAGCTCTATTAGAAGCCGTGGGTCACGGAAGAGAACCATGGAACCCAGTGACTAGTGTTCAGCTCAATTAGGACAAATCTGGGCACTTAGCCATGCAGGAACAATGGCAAGACTTTAGCCCGATCAGGAGCAGCAATGCGTGCCTTGCTAGATCAGGAGCACAGAGGACACCCTGCCGGATCCGAAGGGATGGAAGTCAGCAGCGGGTCTGCGGTGGTGGCAAACAGCAGTGGTGGACAGTGAGCGAAAGCTCAGCTTGAGCCAAGCCATAACAAACATGCACCAGAAGAGAGCGCAGTTGCAAGATTTAATAGAGCGAAAACAGAGCTCCCATACAAAGGGAGGGGACCCAAAGAGGGTAGCCATTGCCAGCTTGAAGGCCTGGGTTTATATCCCGATCATTGTCCCCCTCACTGTGCTCTCAGGCAATAGGTGATTGGCTATTTCTTTACCTCCTGTTTTTGCCTAATTAGCATTTTAGTGAGCTCTCTTTATTACCTGGTTGGTCGGTGTGAGCTAAGTTGCAAGCCCCATGTTTAAAGGTGGATGTGGTCACCTTCCCAGCTAGGCTTAGGGATTCTTAGTCAGCCTAGGAAATCCAGCTTGTCCTGTCTCTCAGTAGCATGAGCAATATTGGTGACAGAGAGGAAGAGCTGAATTCAGTAGACCTGATGATGGTTGGATAG

`>S Yadava-1 18q21.1 W2 gag partial sequence`

GCTCTTCCTCTCTGTCACCAATATTGCTCATGCTACTGAGAGACAGGACAAGCTGGATTTCCTAGGCTGACTAAGAATCCCTAAGCCTAGCTGGGAAGGTGACCACATCCACCTTTAAACATGGGGCTTGCAACTTAGCTCACACCGACCAACCAGGTAATAAAGAGAGCTCACTAAAATGCTAATTAGGCAAAAACAGGAGGTAAAGAAATAGCCAATCACCTATTGCCTGAGAGCACAGTGAGGGGGACAATGATCGGGATATAAACCCAGGCCTTCAAGCTGGCAATGGCTACCCTCTTTGGGTCCCCTCCCTTTGTATGGGAGCTCTGTTTTCGCTCTATTAAATCTTGCAACTGCGCTCTCTTCTGGTGCATGTTTGTTATGGCTTGGCTCAAGCTGAGCTTTCGCTCACTGTCCACCACTGCTGTTTGCCACCACCGCAGACCCGCTGCTGACTTCCATCCCTTCGGATCCGGCAGGGTGTCCTCTGTGCTCCTGATCTAGCAAGGCACGCATTGCTGCTCCTGATCGGGCTAAAGTCTTGCCATTGTTCCTGCATGGCTAAGTGCCCAGATTTGTCCTAATTGAGCTGAACACTAGTCACTGGGTTCCATGGTTCTCTTCCGTGACCCACGGCTTCTAATAGAGCTATAACACTCATCGCATGGTGCAAGATTCCATTCTTTGGAATCCGTGAGGCCAAGGCTTGCCACCATCTTGGAAGTGGCCTGCTGCCATCTTGGGAGCTCTGGGAGCAAGGAACCCCCGGTGACATTTTGGTAACCACGAAGAGATCTCCAAAGCAGTGAGTAATATTGGACCTCTTTCGCTTGCTATTCTGTCCTATCCTTCCTTAGAATTG

`>S Mala-1 4q22.1 H8 env partial sequence`

TGAGAAACATCGCCCCATTATCTCTCCATACCACCCCCAACACTTCAACACTATTTTATTTTTCTTATTAATATAAGAAGACAGGAATGTCAGGCCTCTGAGCCCAAGCCTGCATATATACATCCAGATGGCCTGAAGCAAGTGAAGAATCACAAAAKAAGTGAAAATGGCCGGTTCCTGCCTTAACTGATGACATTACCTTGTGAAATTCCTTCTCCTGGCTCAGAAGCTCCCCCATTGAGCACCTTGTGACCCCTGCCCCTGCCGGCCAGAGAACCCCCTTTGACTGTAATTTTCCATTATCTACCCAAATCCTGTAAAACAGCCCCACCCCTATCTCCCTTYGCTGACTCTCTTTTCTGACTCAGCCCRCCTGCACCCAGGTGATTAAAAAGCTTTATTGCTCACACAAAGCCTGTTTGGTGGTCTCTTCACACGGATGCATGTGAAAATTATACCCCAAGATTTTGCATTATTTTTATCTTGTCTAACTATATAATGAAAACAAATTGAAAAGACCCAATATATTAAAAGCTACTTTATAGGAGTTTTAAAAAGAATTRTATCTATCAAGAATCTCATTATAATTGTAGATACTATGGAGGATTATGGAATAGAGATACACCAAATATTTCCATAATAATAATGTTTTATCTGTTTTTA

`>S Madiga-2 4q22.1 H8 env partial sequence`

CCACTCTAGGTTCCCACACCACCCCCTAATCCCGCTGGAACCAGCCCTGAGAAACATCGCCCCATTATCTCTCCATACCACCCCCAACACTTCAACACTATTTTATTTTTCTTATTAATATAAGAAGACAGGAATGTCAGGCCTCTGAGCCCAAGCCTGCATATATACATCCAGATGGCCTGAAGCAAGTGAAGAATCACAAAAGAAGTGAAAATGGCCGGTTCCTGCCTTAACTGATGACATTACCTTGTGAAATTCCTTCTCCTGGCTCAGAAGCTCCCCCATTGAGCACCTTGTGACCCCTGCCCCTGCCCGCCAGAGAACCCCCTTTGACTGTAATTTTCCATTATCTACCCAAATCCTGTAAAACAGCCCCACCCCTATCTCCCTTCGCTGACTCTCTTTTCTGACTCAGCCCGCCTGCACCCAGGTGATTAAAAAGCTTTATTGCTCACACAAAGCCTGTTTGGTGGTCTCTTCACACGGATGCATGTGAAAATTATACCCCAAGATTTTGCATTATTTTTATCTTGTCTAACTATATAATGAAAACAAATTGAAAAGACCCAATATATTAAAAGCTACTTTATAGGAGTTTTAAAAAGAATTGTATCTATCAAGAATCTCATTATAATTGTAGATACTATGGAGGATTATGGAATAGAGATACACCAAATATTTCCATAATAATAATGTTTTATCTGTTTTTATTTTTAT

`>S Brahmin-1 5p15.31 H2 env partial sequence`

TCTAGGTTCCCATGCCACCCCTAATCCCGCTCAAAGCAGCCTTGAGAAACATTGCTCATTATCTCTCCATACCACCCCCAAAAAATTTTCACCATCCCAACACTTCAGCACTATTTCATTTTTCTTATTAATATAAGAAGACAGGAATATCAGGCCTCTGAGCCCAAGCTAAGCCATCATATCCCCTGTGACCTGCACGTACACATCCAGATGGCCGGTTCCTGCCTTAACTGATGACATTCCACCACAAAAGAAGTGAAAATGGCCTGTTCCTGCCTTAACTGATGACATTGTCTTGTGAAATTCCTTCTCCTGGCTCATCCTGGCTCAAAAGCTCCCCTACTGAGCACCTTGTGACCCCCACTCTGCCTGCCAGAGAACAACCCCCTTTGACTATAATTTTCCTTTATCTACTCACATCCTATAAAATGGCCCCACCCCTATCTCCCTTCGCTGACTCTTTTCGGACTCAGCCTGCCTGCACCCAGGTGATTAAAAGCTTTATTGCTCACACAAAACCTGTTTGATGGTCTCTTCACACGGACGCGCATGAAAGTTACAACTTGAAAATATGATACACACATTAGAAAAAAAGCATGCCAATTAAATGCAATTTATAGAAACACAACAAAAGAAACTCAGAAAAATGATGAAAAAAGATACGGTTCACATACAGCAAGATGGTGTAGCTCCACTCGTGAAACACAATGGACTTTAAAACAACAAGCC

`>S Madiga-2 5p15.31 H2 env partial sequence`

CCCACTCTAGGTTCCCATGCCACCCCTAATCCCGCTCAAAGCAGCCTTGAGAAACATTGCTCATTATCTCTCCATACCACCCCCAAAAAATTTTCACCATCCCAACACTTCAGCACTATTTCATTTTTCTTATTAATATAAGAAGACAGGAATATCAGGCCTCTGAGCCCAAGCTAAGCCATCATATCCCCTGTGACCTGCACGTACACATCCAGATGGCCGGTTCCTGCCTTAACTGATGACATTCCACCACAAAAGAAGTGAAAATGGCCTGTTCCTGCCTTAACTGATGACATTGTCTTGTGAAATTCCTTCTCCTGGCTCATCCTGGCTCAAAAGCTCCCCTACTGAGCACCTTGTGACCCCCACTCTGCCTGCCAGAGAACAACCCCCTTTGACTATAATTTTCCTTTATCTACTCACATCCTATAAAATGGCCCCACCCCTATCTCCCTTCGCTGACTCTTTTCGGACTCAGCCTGCCTGCACCCAGGTGATTAAAAGCTTTATTGCTCACACAAAACCTGTTTGATGGTCTCTTCACAYGGACGCGCATGAAAGTTACAACTTGAAAATATGATACACACATTAGAAAAAAAGCATGCCAATTAAATGCAATTTATAGAAACACAACAAAAGAAACTCAGAAAAATGATGAAAAAAGATACGGTTCACATACAGCAAGATGGTGTAGCTCCACTCGTGAAACACAATGGACTTTAAAACAACAAGC

`>S_Madiga-2_2q34_H4_soloLTR`

TAGGTGAAAGAGCTCACATAGCAGAGTATTGTGATATTAGCCATGATGTACTAACCATCTACATATGTTAGTTGATTGTAATAATAATAAAATAATTAATTTTAAAGATATAAACTATACTTATTGTCAGGTCTCTGAGCCCAAGCCAAGCCATCGCATCCCCTCTGACTTGCAGGTATATGCCCAGATGGCCTGAAGTAACTGAAGAATCACAAAAGAAGTGAAAATGCCCTGCCCCGCCTTAACTGATGACATTCCACCAAAAAAGAAGTGTAAATGGCCGGTCCTTGCCTTAAGTGATGACATTACCTTGTGAAAGTCTTTTTCCTGGCTCATCCTGGCTCAAAAACTCCCCCACTGAGCACCTTGCGACCCCCACTCCTGCCCACCAGAGAACAAACCCCCTTTGACTGTAATTTTCCTTTACCTGACCAAATCTTATAAAACGGCCCCACCCCTATCTCCCTTCTCTGACTCTCTTTTCGGACTCAGCCCGCTTGCACCCAGGTGAAATAAACAGCCATGTTGCTCACACAAAGCCTGTTTGGTCTCTTCACACGGACGTGCATGAAACTTATTAATTACTAAAGATAAATAAGTGAAGTAACACTAGATTCATTTTAACCTGAATTAATACAAATAAGGTGAAATGGTTGGCCTACTGATTGTCTATAAGGAGCAAACCTATAAAATATCAAGTATTGTCACTGCTTGAATATGCAAATGAAAGCTGTGAATAATAATCACGTGGTCAGGGAGAGAAATAAGTTTAAAGATATTAATAAATCTAGAGCACTTTCACATTATGTCAATGTAGTTAAAAGGATGAAAAGAAAATGTTTATCTTCCAGGA

`>S_Luhya-2_3p14.3_H1_soloLTR`

TTCTCTTTGGGAATGTCAGGCCTCTGAGCCCAAGCCAAGCCATCGCATCCCCTATGACATGCACGTACACGCCCAGATGGCCTGAAGTAACTGAAGAATCACAAAAGAAGTGAATATGCCCTGCCCCACCTTAACTGATGACATTCCACCACAAAAGAAGTGTAAATGGCCAGTCCTTGCCTTAACTGATGACATTACCTTGTGAAAGTCCTTTTCCTGGCTCATCCTGGCTCAAAAAGCACCCCCACTGAGCACCTTGCGACCCCCCGCTCCTACCCGCCAGAGAACAAACCCCCTTTGACTGTAATTTTCCTTTACCTACCCAAATCCTATAAAACGGCCCCACCCTTATCTCCCTTCGCTGACTCTCTTTTCCGACTCAGCCCGCCTGCACCCAGGTGAAATAAACAGCCTTGTTGCTCACACAAAGCCTGTTTGGTGGTCTCTTCACACAGACGCGCATGAAAGGGAAGACATACAAAAACAAGGTAAATAAGTAAACTACGTTATATGTTTGATAATGGTGATGTTAAGGGTGGGGAAAGAAGAAAGCAAAGAAGGATAAGAAATGGGAGGGGGCAATTCTAGAAAC

`>S_Luhya-2_5p13.3_K2_soloLTR`

TCCATCATATAAACAGAACTGTGGGGAAAAGCAAGAGAGATCAGATTGTTACTGTGTCTGTGTAGAAAGAAGTAGACATAGGAGACTCCATTTTGTTATGTACTAAGAAAAATTCTTCTGCCTTGAGATTCTGTGACCTTACCCCCAACCCCGTGCTCTCTGAAACATGTGCTGTGTCAACTCAGAGTTGAATGGATTAAGGGCRGTGCAAGATGTGCTTTGTTAAACAGATGCTTGAAGGCAGCATGCTCCTTAAGAGTCATCACCACTCCCTCATCTCAAGTACCCAGGGACACAAAAACTGCGGAAGGCYGCAGGGACCTCTGCCTAGGAAAGCCAGGTATTGTCCAAGGTTTCATAGTCTGAAATATGGCCTCGTGGGAAGGGAAAGACCTGACTGTCCCCCAGCCCGACACCCGTAAAGGGTCTGTGCTGAGGAGGATTAGTATAAGAGGAAGGCATGCCTCTTGCAGTTGAGACAAGAGGAAGGCATCTGTCTCCTGCCTGTCCCTGGGCAATRGAATGTCTCGKTATAAAACCCGATTGTATGCTCCATCTACTGAGATAGGGAAAAACYGCCTTAGGGCTGGAGGTGGGACCTGCGGGCAGCAATACTGCTTTGTAAAGCATTGAGATGTTTATGTGTATGCATATCTAAAAGCACAGCACTTAATCCTTTACATTGTCTATGATGCAAAGACCTTTGTTCACGTGTTTGTCTGCTGACCCTCTCCCACATTGTCTTGTGACCCTGACACATCCCCCTCTTTGAGAAACACCCACGAATGATCAATAAATACTAAKGGAACTCAGAGGCTGGCGGGATCCTCCATATGCTGAACGCTGGTTCCCCGGATCCCCTTATTTCTTTCTCTATACTTTGTCTCTGTGTCTTTTTCTTTCCTAAGTCTCTCATTCCACCTTACGAGAAACACCCACAGGTGTGGAGGGGCAACCCACCCCTACACAGAACCAATGACAAAAACCACATGATTATCTCAATAGATGCAGAAAAGGCCTTCGACAAAATTCAATAGCCATTAATGCTAAAAACTCTCAATAAACTAGGTATCGATGGAACATATCTCAAAATAATAAGCGTTATTTATGACAAACYCACAGCTAATATCATACTGAATGTGCAAAAACTGGAAGCATTCCCTTTGAAAACTGGCAAAAGACAAGTATGCCCTCTCTACCACTATTCAACATAATGTTGTTAGTTCTGGCAAGGGCAATGAGGCCAGAGAAAGAAATAAAGGGTATTTAATTAGGAAAAGAGGAAGTCAAATTGTCCCTGTTTGCAGATGACATGATTGTATATTTAGAAAACCCCATTGTCTCAGCCCAAAATCTCCTTAAGCTGATAAGCAACTTCAGCAAAATCTCAGGATACAAAATCAATGTGCAAAAATCACAACCATTCCTATACATCAATAACAGACAAACAGAGAGCCAAATCATGAGTGAACTCCCATTCACAATTGCTACAAAGTGAATAAAGTACCTAGGAATCCAACTTCCAAGGGATGTGAAGGACCTCTTCAAGGAGAACTACAAACCACTGCTCAACAAAATAAAAGAGGACACAAACAAATGGAAGAATATTCCATGCTCATGGATAGGAAGAATCAATATTGTGAAAATGGCTATACAGGTCAAGGTAATTTATAGATTCAATGCCATCCCCATCAAGCTACCAATGACTTTCTTCACAGAATTGGGAAAACTACTTTAAAGTTCATATGGAACCAAAAAA

